# Face familiarity detection with complex synapses

**DOI:** 10.1101/854059

**Authors:** Li Ji-An, Fabio Stefanini, Marcus K. Benna, Stefano Fusi

## Abstract

Synaptic plasticity is a complex phenomenon involving multiple biochemical processes that operate on different timescales. We recently showed that this complexity can greatly increase the memory capacity of neural networks when the variables that characterize the synaptic dynamics have limited precision, as in biological systems. These types of complex synapses have been tested mostly on simple memory retrieval problems involving random and uncorrelated patterns. Here we turn to a real-world problem, face familiarity detection, and we show that also in this case it is possible to take advantage of synaptic complexity to store in memory a large number of faces that can be recognized at a later time. In particular, we show that the familiarity memory capacity of a system with complex synapses grows almost linearly with the number of the synapses and quadratically with the number of neurons. Complex synapses are superior to simple ones, which are characterized by a single variable, even when the total number of dynamical variables is matched. We further show that complex and simple synapses have distinct signatures that are testable in proposed experiments. Our results indicate that a memory system with complex synapses can be used in real-world tasks such as face familiarity detection.

**Significance:** The complexity of biological synapses is probably important for enabling us to remember the past for a long time and rapidly store new memories. The advantage of complex synapses in terms of memory capacity is significant when the variables that characterize the synaptic dynamics have limited precision. This advantage has been estimated under the simplifying assumption that the memories to be stored are random and uncorrelated. Here we show that synaptic complexity is important also in a more challenging and realistic face familiarity detection task. We built a simple neural circuit that can report whether a face has been previously seen or not. This circuit incorporates complex synapses that operate on multiple timescales. The memory performance of this circuit is significantly higher than in the case in which synapses are simple, indicating that the complexity of biological synapses can be important also in real-world memory tasks.

## Introduction

Synaptic memory is a complex phenomenon, which involves intricate networks of diverse biochemical processes that operate on different timescales. We recently showed that this complexity can be harnessed to greatly increase the memory capacity^1, 2^ in situations in which the synaptic weights are stored with limited precision. More specifically, we proposed a complex synaptic model in which *m* variables that might correspond to different biochemical processes interact within each synapse such that the memory capacity of a population of synapses, estimated by an ideal observer who has access to all the synaptic weights, can increase almost linearly with its size (i.e., the number of synapses *N_syn_*), even when both *m* and the number of states of each variable grow no faster than logarithmically with *N_syn_*. This is the optimal scaling under some conditions (see ^2^) and significantly better than what can be achieved by employing a simple synapse characterized by a single variable^3–5^.

These previous studies on complex synapses focused on the problem of storing a large number of random and uncorrelated memories. Only recently, complex synapses started to be employed in more realistic problems (e.g., see^6^) in which memories are structured and correlated. Here we show that synaptic complexity can be important also in a real-world problem, face familiarity detection. The task is particularly difficult because we require that each face is presented only once (one-shot learning) and it has to remain recognizable for a long time. This is a typical situation in which complexity can play an important role. Indeed, the complex synapses of ^2^ that we incorporated into our model are characterized by dynamical variables that operate on multiple timescales. The fast ones can rapidly store information about a new visual stimulus such as a face, even when the stimulus is shown only once. This information is then progressively transferred to the slow variables, which can retain it for a long time. Because of these slow variables, which influence the synaptic efficacy, the older memories are protected from overwriting due to the storage of new faces. Synapses that are described by a single dynamical variable can either learn quickly if they are fast, but then they also forget quickly, or they can retain memories for a long time if they are slow, but then they cannot learn in one shot and require multiple exposures to the same face. This plasticity-rigidity dilemma concerns a very broad class of realistic synaptic models whose dynamical variables have a limited precision^3, 5, 7^.

Our memory benchmark for the complex synapses, familiarity detection (sometimes called familiarity discrimination or novelty detection) is an important component of recognition memory, which has been widely studied in humans and in animals. In particular, familiarity detection refers to the ability to rapidly memorize new items and report at a later time whether we have encountered them or not. In the case of faces, we would report that a face of a person is familiar if we experience the sense that we have already encountered that person in the past. The second component of recognition memory is recollection, which corresponds to the retrieval of the details of the individual (e.g. the name) and the episodic memories associated with that person. We can often experience a sense of familiarity without being able to recollect the details about an encountered individual. Familiarity detection, which is the focus of this article, has been studied in the famous and remarkable experiment by Standing ^8^, in which he showed that it is possible to recognize a surprisingly large proportion of 10,000 images that are flashed on a screen only once and for a brief time. The subjects were asked whether they had seen an image or not, which is one way of assessing the familiarity of an image. Although familiarity detection is only one component of recognition memory, in the article we will use the verb ’recognize’ to indicate the ability of a subject to report whether a visual stimulus had already been seen or not. The result of the Standing experiment is even more remarkable when one considers that more recent studies proved that subjects could memorize many details about each image^9,10^. The neural substrate of recognition memory is unknown, although multiple lesion studies indicate that the hippocampus and perirhinal cortex play an important role^11–13^. The role of each area is controversial as for some investigators both the hippocampus and the perirhinal cortex contribute to recollection (memory retrieval) and familiarity^13, 14^ and for others the hippocampus supports recollection only, and perirhinal supports familiarity ^11, 12^. One of the problems in the interpretation of these studies is that it is difficult to separate the contribution that each area gives to familiarity and recollection because when a memory can be recollected it can always be recognized. Another problem is that the role of these two areas differs depending on the nature of the memories (e.g. recognition of novel faces is intact in patients with lesioned hippocampus at a short retention interval, instead recognition memory for words, buildings, inverted faces, and famous faces is impaired^15^), on the length of the retention interval (for intervals of a few minutes or longer the hippocampus is certainly important for familiarity ^13, 16^) and on whether the memory is presented in a particular context or in isolation (perirhinal cortex is more important for the recognition of items in isolation whereas the hippocampus is more important when there is a contextual or associational component^11^). In the Discussion we will describe a possible interpretation of our model.

There are several biology-inspired computational models studying different aspects of recognition memory: some neural network models following the complementary learning systems approach were proposed to tease apart the hippocampal and neocortical contributions to recognition memory^17, 18^; other models were concerned with the synaptic plasticity (learning) rules in the perirhinal cortex^19^. Finally, there are models that stress the distinct roles for familiarity and recollection in retrieving memories^20^.

Analytical estimates of familiarity memory capacity showed that in the case of random uncorrelated patterns, the number of memories that can be correctly recognized as familiar can scale quadratically with the number of neurons *N* in a recurrent network^21^. Not too surprisingly, this is a much better scaling than the linear scaling of the Hopfield model ^22^, in which random memories are actually reconstructed (see also the Discussion). The scaling for memory reconstruction is markedly worse and can be as low as 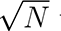 when the patterns representing the memories are correlated^19^. These computational models can replicate some interesting aspects of experiments on the capacity of human recognition memory^23^.

We constructed a model for recognition memory that, for the first time, incorporates complex synapses characterized by variables that have limited dynamical range (number of distinguishable states). We show that a simple neural circuit designed to reconstruct the memorized face can take advantage of the complexity of synapses and can efficiently store a large number of faces. In particular, we show that the number of faces that can be successfully recognized as familiar scales approximately quadratically with the number of neurons, or linearly with the number of synapses. This is the same scaling achieved in^21^, in which synaptic weights could be stored with unlimited precision. Moreover, this scaling is similar to the one predicted for random patterns in^2^. Interestingly, the network can recognize a face even when it is presented in a different pose, and the scaling is only slightly worse than in the case in which the exact same picture of the face is presented for familiarity testing. This ability to generalize is a distinctive feature of recognition memory, it is observed in experiments and it plays an essential role in any machine learning system that relies on novelty signals to speed up learning^24^. We then compared the performance of the recognition system with complex synapses to one with the same architecture, but with a larger number of neurons and simple synapses characterized by a single dynamical variable. The number of neurons is chosen so that the total number of synaptic variables would be the same in the two systems. We show that the system with complex synapses outperforms the one with simple synapses, indicating that complexity provides the neural system with a clear computational advantage.

## Materials and methods

### Face data set

We used a large-scale face data set called VGGFace2^25^. Compared to other public face data sets (such as Labelled Faces in the Wild data set^26^, CelebFaces+ data set^27^, VGGFace data set^28^, MegaFace data set^29^, and Ms-Celeb-1M data set^30^), it contains a relative large number of individuals (3.31 million images of 9131 individuals) and large intra-identity variations in pose, age, illumination and background (362.6 images per person on average), with available human-verified bounding boxes around faces. For each face image, the bounding box was then enlarged by 30% to include the whole head, resized such that the shorter side was 256 pixels long, and center-cropped to 224 *×* 224 pixels to serve as the input for our neural system described below.

### Neural face familiarity detection system

Our face familiarity detection system consists of three modules: an input (embedding) module, a memory module, and a readout (detection) module.

#### Input (embedding) module

The embedding module consists of a deep convolutional neural network (SE-ResNet-50, SENet for short), which is a ResNet architecture integrated with Squeeze-and-Excitation (SE) blocks adaptively recalibrating channel-wise feature responses^31^. Such networks for face recognition with different architectures and different training protocols are publicly available online^25^. We used one specific version of SENet, which is pre-trained on the MS-Celeb-1M data set^30^ and then fine-tuned on the VGGFace2 data set. This version was reported to have the best generalization power on face verification and identification among architectures (e.g., SENet and ResNet-50) and training protocols (e.g., training on different data sets with or without fine-tuning) tested^25^.

The 2048 dimensional activity of the penultimate layer (adjacent to the classification layer) was extracted as the face feature vector for each face image input. Because the face feature vectors are sparse and non-negative, we took the following steps to transform them into a format that’s suitable as the input to the memory module: (i) the dimensionality of the feature vector of each face was first reduced using principal component analysis (PCA); (ii) each dimension was then binarized with a threshold equal to the median (−1 for values less than the median and +1 for values larger than the median). The first *N* binarized principal components were taken as the binary face pattern *x* = [*x*_1_*, . . . , x_N_*]*^T^*, serving as the activity of the *N* input neurons of the memory module.

The data set contains faces from 9131 different people. To facilitate the evaluation of the memory performance of our system over a vast time scale, a larger number of independent non-evaluated patterns are required to be stored in between the face patterns whose memory signals are being tracked. Thus, we synthesized artificial face patterns matching the first and second moments of real face patterns. First, we extracted the mean values and the diagonal covariance matrix of the face feature vectors after PCA to get an estimate of the distribution of patterns generated by the faces of all the people in the data set. We then synthesized artificial face patterns by passing new samples from the corresponding multivariate normal distribution through the binarization step. Mathematically, this process is equivalent to generating unstructured, random, binary patterns. These artificial patterns were presented to the neural system to be memorized at time steps in between the storage of the real face patterns, but were not used to evaluate the memory performance.

#### Memory module

The memory module is the only part of our network containing plastic synapses. The synapses are continuously updated by the ongoing presentation of the face patterns, whereas the weights of the input module are frozen during the online learning phase. The memory module consists of *N* memory neurons, one for each unit of the embedding module providing inputs to the memory module. We will refer to these units as input neurons for short. The *j*-th input neuron connects to the *i*-th memory neuron (for *i* ≠ *j*) with synaptic weight (efficacy) *w_ij_* and bias term *b_i_*. There is no connection between the *i*-th input neuron and the *i*-th memory neuron for any *i* (i.e., *w_ii_* = 0 *∀i*). The activity of the *i*-th memory neuron is

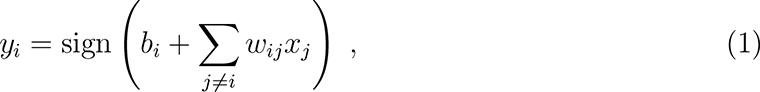

and we denote the binary memory patterns retrieved in this manner as *y* = [*y*_1_*, . . . , y_N_*]*^T^* . This plastic layer of synapses implements a simple feedforward memory model that can perform an approximate one-step reconstruction of a stored input pattern from a noisy cue at test time. Because the *i*-th memory neuron *y_i_* is expected to reconstruct the *i*-th input neuron *x_i_*, we set the value of the *i*-th memory neuron to be *x_i_* during learning.

To update the synaptic weights and biases we used the learning rule:

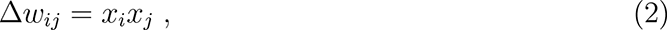

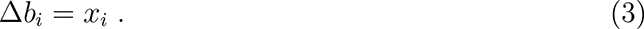

These equations describe the desirable plasticity steps to store each new pattern. However, simply applying these additive updates would eventually result in unbounded values of the *w_ij_*. Therefore, we employed a mechanism to limit the weights to bounded dynamical ranges. For each synapse (i.e., for each weight *w* and bias term *b*), we implemented a complex synaptic model^2^ with *m* dynamical variables *u*_1_*, . . . , u_m_* in discrete time. Here *m* denotes the total number of variables per synapse (a measure of synaptic complexity), each of which operates on a different timescale. Specifically, at each time step *t* the dynamical variables *u_k_* (for 2 *≤ k ≤ m*) are updated as follows (the indices *i* and *j* labeling the synapses are omitted for simplicity)

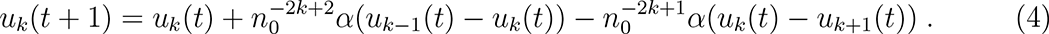

For *k* = *m*, the last variable *u_k_*_+1_ is simply set to zero in this update equation, and for *k* = 1 we have

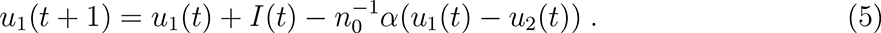

Here *I*(*t*) is the desirable update (Δ*w* or Δ*b*) imposed by the pattern *x*(*t*), which takes a value +1 or *−*1 and is computed from equations (2) or (3). The first variable *u*_1_ is used as the actual value of the synaptic weight *w* or bias *b* at test time. The parameters *α* and *n*_0_ determine the overall timescale of the model dynamics and the ratio of timescales of successive synaptic variables (we set *α* = 0.25 and *n*_0_ = 2 in our models; see^2^ for additional details).

To study the situation in which variables can only be stored with limited precision, we discretized the *m* synaptic variables and truncated their dynamical range to a maximum and minimum value. Hence, each variable can take one of only a finite number of integer-spaced values arranged symmetrically around zero (namely *{−V, −V* + 1*, . . . , V −* 1*, V }*, where in our simulations we chose *V* = 31*/*2), corresponding to 32 levels (5 bits). At every time step, if the *u_k_*(*t* + 1) computed according to equations (4) and (5) falls between two adjacent levels, its new value is set to one of those two levels, based on the result of a biased coin flip with an odds ratio equal to the inverse ratio of the distances from *u_k_*(*t* + 1) to the two levels.

#### Readout (detection) module

The readout (detection) module compares the output *x* = [*x*_1_*, . . . , x_N_*]*^T^* of the embedding module and the output *y* = [*y*_1_*, . . . , y_N_*]*^T^* of the memory module to assess the level of familiarity of a given pattern. This module computes the Hamming distance between *x* and *y*, and outputs “familiar” (or “unseen”/“unfamiliar”/“novel”) if the distance is smaller (or larger) than some pre-set threshold. This approach is similar to the one proposed in^21^.

### Evaluating the memory signal and noise

One way to measure the strength of a memory is to take the perspective of an ideal observer, who can access directly all the synaptic weights^1, 2, 7^. Following this approach, we considered the expected ideal observer signal *S*_io_ and noise term *N*_io_, and computed the ideal observer signal-to-noise ratio (ioSNR) *S*_io_*/N*_io_(Δ*t*) as a measure of the expected fidelity of recall of the stored memories as a function of the time elapsed since storage. The *S*_io_ and *N*_io_ are computed as follows.

For a given face memory, the signal at time *t* of the input pattern *x*(*t^t^*) stored at an earlier time *t^t^*is defined as the overlap (inner product) between the synaptic modification Δ*w_ij_*(*t^t^*) imposed at storage and the current ensemble of synaptic weights *w_ij_*(*t*):

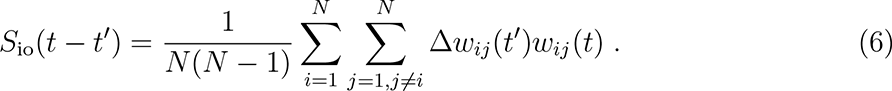

We can then compute the average (denoted by ()) over all memories with an age of Δ*t* = *t−t^t^* to obtain the expected signal

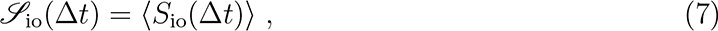

and the corresponding noise term

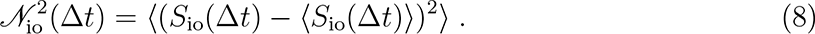

We also considered another measure of the memory signal that is more directly related to the ability of the system to read out the stored memory, the readout signal *S*_r_. Similarly to the ioSNR, this signal is defined as the overlap between an input pattern *x*(*t^t^*) (stored at time *t^t^*) and the retrieved memory pattern *y*(*t, t^t^*), the output of the memory module when the same pattern is presented again at time *t* without updating the synaptic weights. We have

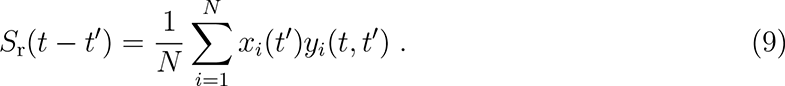

As above, we can compute the expected signal *S*_r_ and noise *N*_r_ by averaging over memories of a given age, and obtain the readout signal-to-noise ratio (rSNR) *S*_r_*/N*_r_(Δ*t*).

Because the SNR decreases with time, we define the quantities *t^∗^*_ioSNR_ and *t^∗^*_rSNR_ as the memory ages *t^∗^* at which the ioSNR or rSNR, respectively, drop below a certain retrieval threshold as measures of the familiarity memory capacity of the system or of the expected memory lifetime.

### Task protocol

To evaluate the performance of our system, we considered two tasks in which we presented a series of pre-processed face images to the neural system and tested its memory on randomly chosen faces (see Fig. 1c). We considered two types of tests. In the familiarity detection (FD) test, the neural system is required to determine whether the face image presented at test time is familiar or unseen by comparing the output of the detection module to a threshold. A familiar face image could be a previously stored one (same pose, SP) or a new pose of a previously presented face (different pose, DP), while an unseen face image is an image of an unseen person. In the two-alternative forced-choice (FC) task, the neural system is presented with a pair of face images containing one familiar (either SP or DP) and one unseen face, and is required to choose which one of the two is familiar by comparing the output of the detection module for the two faces. These tasks are made particularly challenging by the fact that the familiar faces are presented only once.

**Figure 1:**
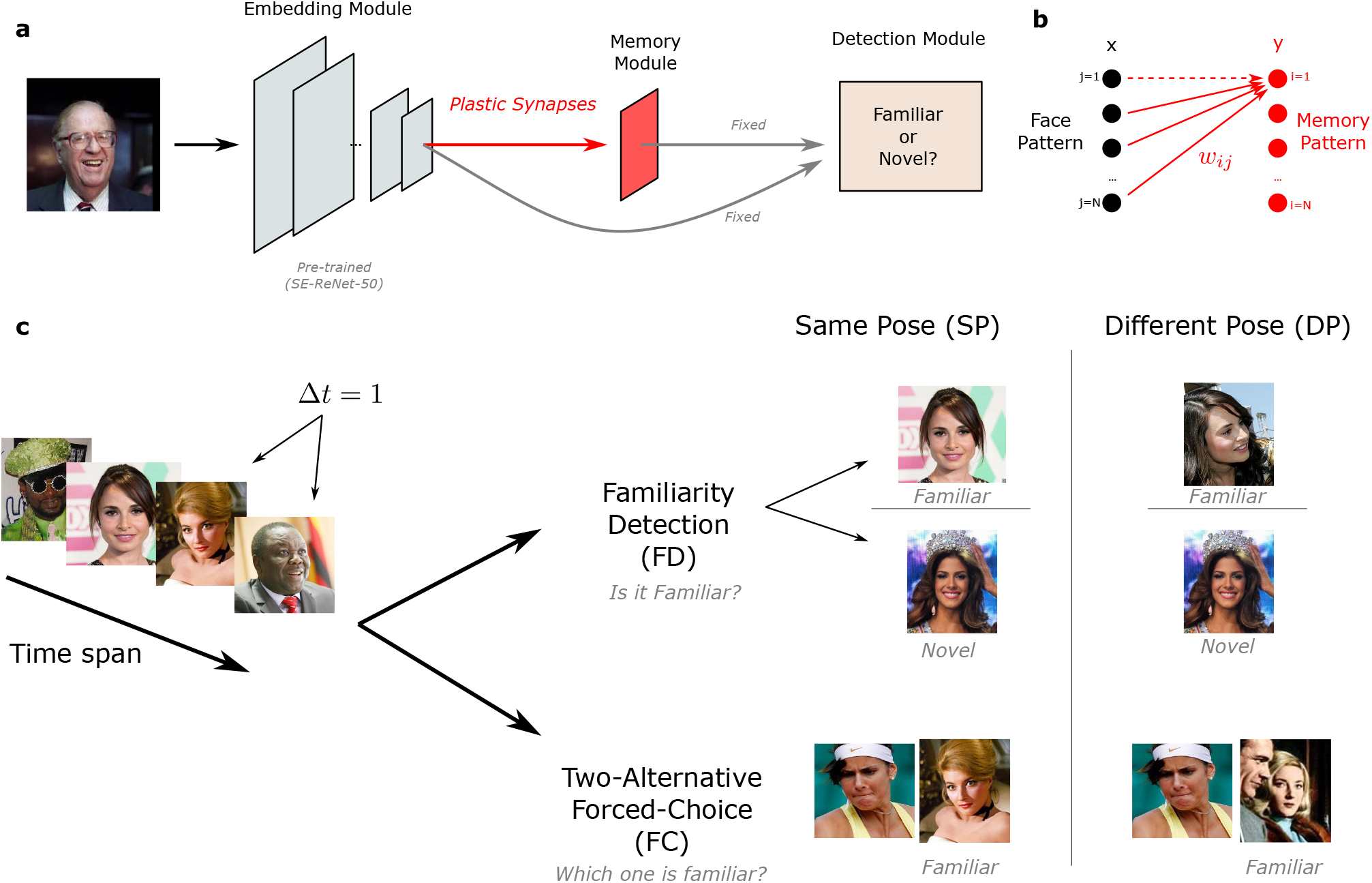
The architecture of our face familiarity detection system and the task diagram. (a) The neural system contains three modules: the input (embedding) module, the memory module, and the readout (detection) module. The synapses between the embedding module and the memory module (as well as the biases in the memory module) are plastic, while all other synapses are fixed (after being either set by hand or pre-trained) during the test phase, which requires online learning of face patterns. (b) The plastic connections between the input neurons in the embedding module and the memory neurons in the memory module. (c) A series of face images are presented to the neural system. In each familiarity detection (FD) test, the system is required to determine whether a presented face is familiar or unseen. A face is considered familiar if the test image is identical to a previously presented one (i.e., the same pose, SP) or a new pose of a previously presented face (i.e., a different pose, DP), and is considered novel if it is an image of an unseen person’s face. In each two-alternative forced-choice (FC) test, the neural system is presented with a pair of face images (exactly one familiar and one unseen), and is required to choose which one of the two is familiar.

### Evaluating the task performance

In the FD test, the face images presented to the system are balanced, i.e., familiar faces previously presented within a certain age-range and unseen faces appear at test time with equal probability. The threshold on the overlap in each model was optimized for face memories within the model-specific age-range, which was operationally set to be the memory lifetime *t^∗^*_ioSNR_. We then computed the task classification performance for face memories of different ages as a function of the time elapsed since storage (also see Appendix I).

In the FC test, the task performance is defined as the probability of correctly choosing the familiar face (over the unseen one) for face memories of different ages. Similarly to our SNR analyses, we defined the memory age *t^∗^* as the age at which the task performance drops below some threshold (which defines the quantities *t^∗^*_FD_ and *t^∗^*_FC_).

In each simulation, all memory quantities, including the SNR and the task performance, are evaluated after the neural system reaches its steady state, i.e., when a large number of face patterns (with constant input statistics) have already been stored. In the steady state, the distribution of synaptic weights does not change any longer^2^, although synapses continue to be updated as new face images are memorized. The system is then presented with two thousand real face images from different people, followed by the necessary number of synthesized face image patterns. Our memory measures were evaluated only over these real face images and further averaged over independent simulations to reduce the noise floor.

### Advanced learning schedules and idealized decay models

In addition to the simple schedule described above in which each face is stored only once, we further consider two types of advanced learning schedules (namely optimal and pre-determined learning schedules), where the same image pattern can be presented more than once and is evaluated during each presentation.

Under the optimal learning schedules, the ideal observer signal (ioSignal) of a specific pattern is constantly monitored after its initial presentation. Every time its ioSignal drops below the pre-specified threshold, the same pattern is presented again (refreshed) to boost its memory strength. The length of the *n*-th interval between the *n*-th and (*n* + 1)-th presentations will vary with the interval number *n*. Even though monitoring the memory signal in real time (without modifying it) is not feasible in experiments, we will show that this theoretical analysis of the length of the *n*-th interval reveals major differences between synaptic models with different complexity, which have measurable consequences.

Motivated by theoretical results on optimal learning schedules, we also propose pre-determined learning schedules, under which the length of the interval between each two consecutive presentations takes the form of *γn^β^*, where *n* is the interval number, *γ* is the length of the first interval (between the first and the second presentations), and *β* is the exponent. When *β* takes value 0 or 1, this schedule degenerates into constant or linear schedules, in which a specific pattern is presented again (refreshed) after a interval of a constant length or after a interval of a linearly increasing length.

To facilitate the study of the behavior of our simulated models with simple and complex synapses under these schedules, we introduced three idealized decay models in which the memory signal decays with a pre-specified profile: exponential, inverse-square-root power-law, and hyperbolic power-law. This allows us to quickly run many numerical experiments with different learning schedules, since we do not have to simulate the internal dynamics of the complex synapses, which is inherently stochastic and would require averaging over many realizations to obtain an estimate of the expected behavior (which is instead represented directly by the specified decay function). These idealized models will allow us to demonstrate that pre-determined learning schedules can be used in experiments to discriminate different decay profiles, which are related to the complexity of a memory system, without the need to access its internal constituents.

Free parameters in the idealized decay models were chosen to match the behavior of the simulated models with simple or complex synapses:

1. The exponential decay model, with memory signal *r*(*t*) = *C*_exp_*e^−t/τ^*^exp^, where *C*_exp_ = 1 is the initial memory strength and *τ*_exp_ = 7.486 is the time constant, fit to the signal strength of simulated models with simple synapses (*m* = 1).
2. The inverse-square-root power-law decay model, with 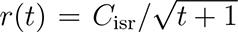, where *C*_isr_ = 1.316 is the initial memory strength in order to fit the power-law decay regime of signal strength of models with complex synapses (*m* = 8).
3. The hyperbolic power-law decay model, with *r*(*t*) = *C*_hyp_*/*(*t* + 1), where *C*_hyp_ is equal to *C*_isr_.

## Results

As we were interested in the scaling properties of the familiarity memory capacity in the case of familiarity detection, we systematically studied the performance of our neural system as functions of two key parameters: the number of memory neurons *N* and the number of dynamical variables *m* per synapse (i.e., the synaptic complexity). Increasing the synaptic complexity *m* for constant *N* or increasing *N* for constant *m* initially leads to a rapid improvement of the familiarity memory capacity but only up to a point where *N* is comparable to the longest timescale of the synapse determined by *m*. Beyond this point, the growth of the familiarity memory capacity slows down (and it may even drop slightly in the case of increasing *m* further). To take advantage of a larger population of neurons, it is important to increase the longest timescale of the synapses, which is related to its complexity *m*. This can be achieved by choosing an *m* that grows logarithmically with *N* (such that *m* = log_2_ *N −* 1, as suggested in^2^). We present the results of varying *m* and *N* separately in Appendix II.

### SNR analysis and memory performance

We considered the SP and the DP tests separately (see Fig. 2). The DP performance cannot surpass the SP performance, because detecting familiarity for a different pose of the same person is clearly more difficult.

**Figure 2:**
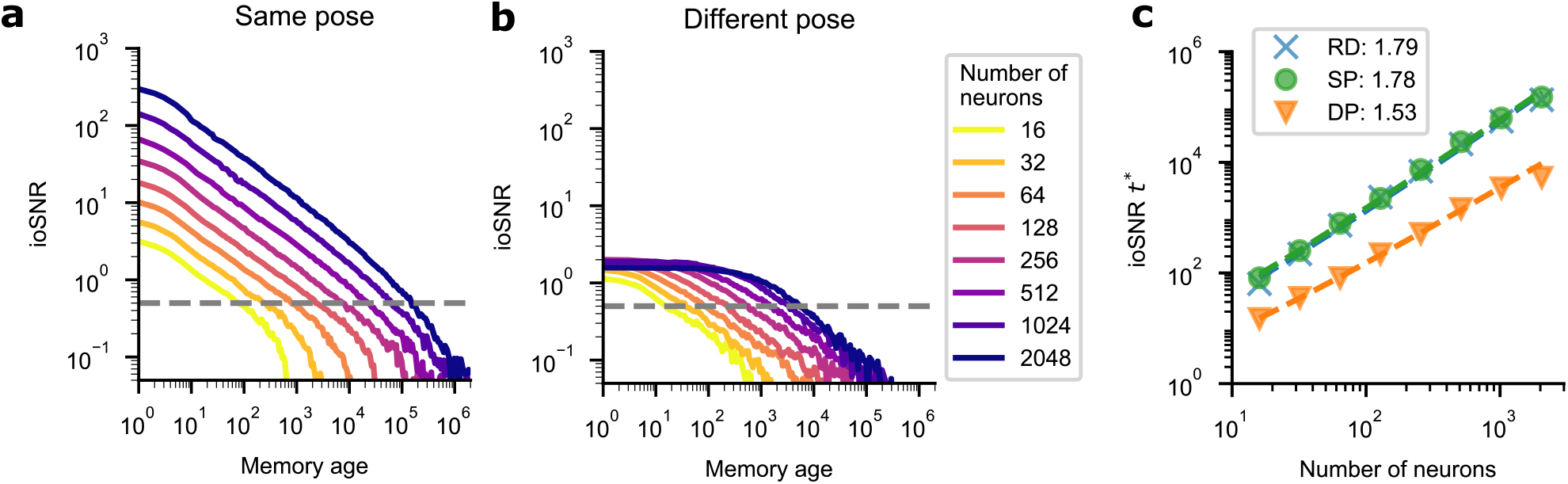
Ideal observer SNR (ioSNR) of the memory module as a function of face memory age and its scaling properties. (a) Doubly logarithmic plots of ioSNR versus the number of subsequently stored memories. Different curves correspond to models with a different number *N* of memory neurons and *m* of dynamical variables in the same pose (SP) case. The parameters *N* and *m* are varied by increasing *N* by factors of two and setting *m* = log_2_ *N* 1. (b) The same as in the previous panel, but in the different pose (DP) case. (c) Log-Log plot of the ioSNR memory lifetime versus *N* in the SP, DP, and random-pattern (RD) cases. The legend indicates the best fit linear regression slopes (corresponding to the power of base *N* in the scaling behavior with the logarithmic correction; data points deviating from the regression line due to saturation were excluded).

The ioSNR critically depends on the number of memories that are stored after the tracked face pattern, i.e., the memory age (see Materials and methods). Different curves correspond to synaptic models with different numbers of input features (and memory neurons) *N* and dynamical variables *m*. The curves are plotted on a log-log scale, for which a straight line represents a power-law dependence.

In the SP case, the ioSNR curves decay as a power-law over a time interval *T* corresponding to the longest timescale of the synapse before the decay becomes exponential. The ioSNR decays as slowly as the inverse square root of the memory age in the power-law regime. Changing *N* shifts the ioSNR curves in the log-log plot vertically, while increasing *m* primarily extends the power-law regime (i.e., increases *T* ; see Fig. 2a). We determined the scaling of the familiarity memory lifetime with *N* (and *m*), where the lifetime *t^∗^*_ioSNR_ is represented by the memory age at which the ioSNR first drops below a given threshold. A value of 1 corresponds to a situation where the signal and the noise are of the same intensity. We chose a threshold of 0.5, though its precise value does not affect the scaling behavior much. We found that the familiarity memory lifetime scales approximately as *N*^2^ (see Fig. 2c, in which the linear regression slope on a log-log scale is about 1.78 for the SP case, compared to 1.79 for random patterns (RD)). This scaling is very close to the theoretical result for optimal storage of random unstructured patterns^2^. Because *m* increases together with *N* (logarithmically), the familiarity memory lifetime scales exponentially with *m* (with the same linear regression slope on a log_2_-linear plot of *t^∗^*_ioSNR_ versus *m*).

For the DP case (see Fig. 2b), the ioSNR curves are lower than those in the SP case, due to the differences between the memorized and the tested face patterns. When there are more memory neurons, the shape of its initial decay with memory age becomes flatter. The initial ioSNR increases slowly with *N* for *N <* 512, and then drops a little for larger *N*, because compared with the former features, latter features are much less correlated between poses of the same person. Nevertheless, the familiarity memory capacity still scales as a power of *N* : the regression slope is 1.53 (the model with 2048 neurons is removed from linear regression due to saturation).

We also studied the properties of the rSNR. We found that the rSNR behaves similarly to the ioSNR at long time lags, but deviates from it for small memory ages, reflecting the effect of the neuronal nonlinearity. This nonlinear effect, which becomes more significant for larger *N* or smaller *m* (see also Fig. 5), leads to larger initial rSNR values, but does not substantially affect the memory lifetime compared to the ioSNR measure. The initial SNR enhancement quickly attenuates, leading to a similar scaling for the familiarity memory lifetime *t^∗^*_rSNR_. These results further validate the ideal observer approach.

### Task performance

In Fig. 4, we plot the task performance in the FD test as a function of memory age. In the SP case, increasing *N* and *m* leads to a substantial extension of the task-relevant familiarity memory lifetime *t^∗^*_FD_ (see Fig. 4a). The memory lifetime was estimated assuming a performance threshold of 60% (this value was chosen to keep the initial task performance of all the simulations above the threshold). The power-law scaling behavior of the familiarity memory lifetime is revealed by plotting *t^∗^*_FD_ versus *N* on a log-log scale (linear regression slope 1.78; see Fig. 4e), which shows a very similar growth also in the RD case (linear regression slope 1.85). The initial task performance cannot reach 100% because each model optimized for the model-specific age-range has a non-zero constant error rate for unseen faces, even the true positive rate (accuracy for familiar faces) saturates at 100% (also see Appendix I).

**Figure 3:**
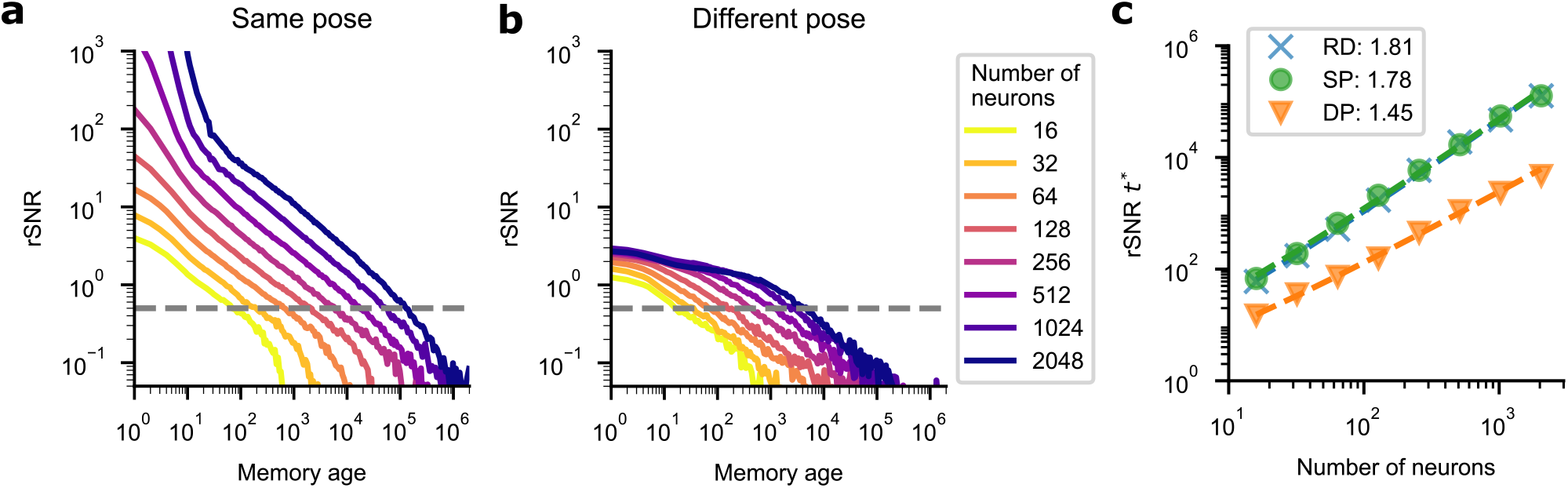
Readout SNR (rSNR) of the memory module as a function of face memory age and its scaling properties. (a) Doubly logarithmic plots of rSNR versus the number of subsequently stored memories. Different curves correspond to models with a different number *N* of memory neurons and *m* of dynamical variables in the same pose (SP) case. The parameters *N* and *m* are varied by increasing *N* by factors of two and setting *m* = log_2_ *N* − 1. (b) The same as in the previous panel, but in the different pose (DP) case. (c) Log-Log plot of the rSNR memory lifetime versus *N* in the SP, DP, and random-pattern (RD) cases. The legend indicates the best fit linear regression slopes (corresponding to the power of base *N* in the scaling behavior with the logarithmic correction; data points deviating from the regression line due to saturation were excluded).

**Figure 4:**
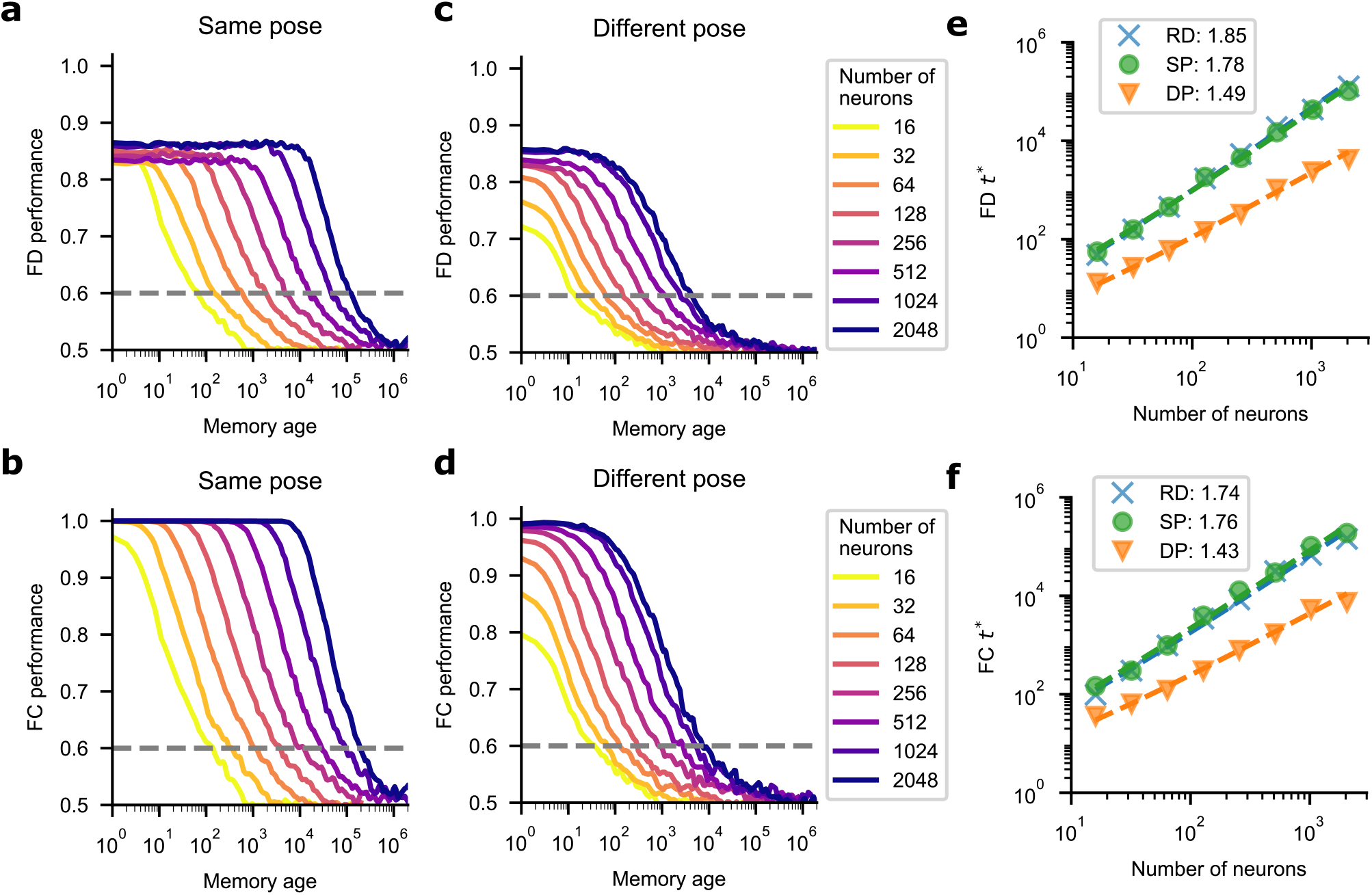
Familiarity detection (FD) and two-alternative forced-choice (FC) test performance of our system and their scaling properties. (a, b) Task performance as a function of the memory age. Different curves correspond to models with a different number *N* of memory neurons (and number *m* of dynamical variables such that *m* = log_2_ *N* 1) in the same pose (SP) case. (c, d) As in the previous panels, but for the different pose (DP) case. (e, f) FD and FC memory lifetimes versus *N* in the SP, DP, and random-pattern (RD) cases. The legend indicates the best fit linear regression slopes (corresponding to the power of base *N* in the scaling behavior with the logarithmic correction; data points deviating from the regression line due to saturation were excluded).

As expected, in the DP case the task performance is worse than in the SP case (see Fig. 4c). However, we still found a reasonable power-law scaling with *N* (regression slope 1.49).

We also plotted the task performance in the FC test as a function of the memory age. The regression slope of the memory lifetime *t^∗^*_FC_ versus *N* on a log-log scale is 1.76 for the SP case and 1.43 for the DP case (see Fig. 4b, d and f).

### Complex versus simple synapses

To obtain a fair comparison between complex synapses ^2^ and the well-studied, simple (multi-state) synapses^3, 4, 32^, we evaluated the familiarity memory performance of a neural system with complex synapses and three models with simple synapses with an approximately equal number of dynamical variables. The complex model has 512 memory neurons, and every neuron has 511 incoming synapses and one bias with eight dynamical variables each (i.e., 512^2^ *∗* 8 = 2097152 variables in total). All of the three simple models have 1448 memory neurons, and every neuron has 1447 incoming synapses and one bias with one dynamical variable each (i.e., 1448^2^ *∗* 1 = 2096704 variables in total). These simple synapses follow essentially the same model dynamics as the previously studied hard-bounded multi-state synapses^4^. They differ in their level of plasticity: the synapses in the first model are updated every time an input pattern is stored, while the synapses in the second and third ones are changed stochastically according to a learning rate (encoding probability) less than one, and thus are more “rigid”^3, 33^. Small learning rates lead to lower initial ioSNR values, but also to longer memory lifetimes. We choose *q* = 0.128 for the second model so that its initial ioSNR is comparable to the complex synapse system in the SP and DP cases. For the third model, we picked *q* = 0.01 to obtain the longest memory lifetimes possible for a system of simple synapses of this size, with an initial SNR just above the threshold (in the DP case).

Each variable in all of these models has the same number of discrete levels, and the total numbers of variables are approximately matched in the simple and the complex system. These simulations show that the complex system has a substantially better familiarity memory performance than the simpler systems (see Fig. 5), despite the smaller number of neurons. For the SP and RD cases, the memory lifetime of the system with complex synapses is *∼* 300 *−* 500 times longer; while for the DP case, the improvement factor is *∼* 30 *−* 60. Slower simple synapses (with smaller *q*) can greatly extend the familiarity memory lifetime, but at the expense of the initial SNR and thus the generalization ability. Even so, they are far from matching the memory lifetime of the complex system. We can conclude that the memory model with complex synapses performs at least two orders of magnitude better in terms of familiarity memory capacity, and we expect the gap between simple and complex systems to grow even wider in networks with a larger number of neurons because of the different scaling behaviors.

**Figure 5:**
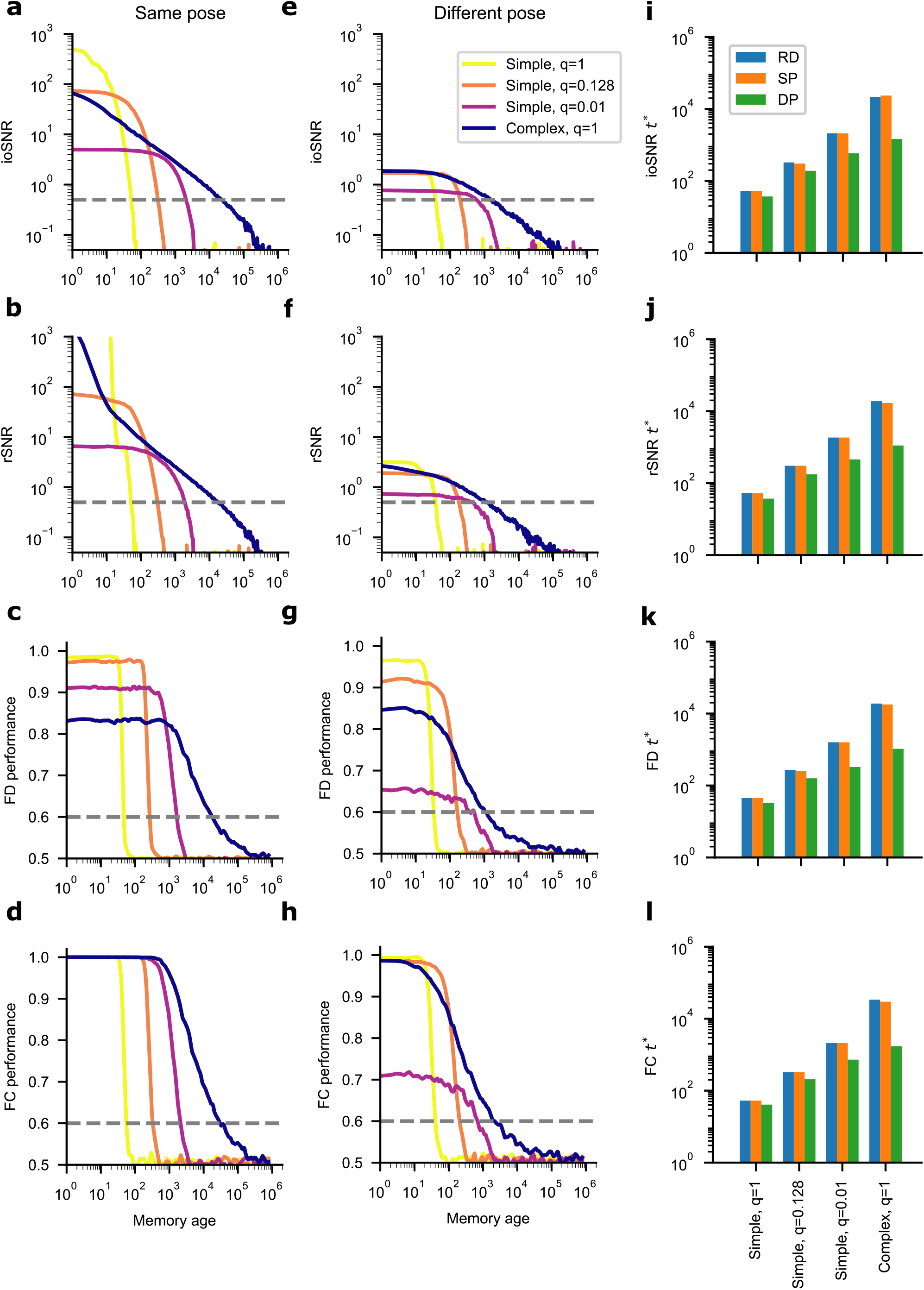
Comparison between models with simple synapses (*N* = 1448, *m* = 1) and different learning rates (*q* = 1, 0.128, and 0.01, respectively) and a complex model (*N* = 512, *m* = 8, *q* = 1) with approximately the same total number of plastic variables. (a-d) Comparisons between models for the same pose (SP) case in terms of ioSNR, rSNR, familiarity detection (FD) performance, and two-alternative forced-choice (FC) performance. (e-h) Similar comparisons between models in the different pose (DP) case. (i-l) Comparisons between models in terms of different measures of familiarity memory lifetime (*t^∗^*_ioSNR_, *t^∗^*_rSNR_, *t^∗^*_FD_, and *t^∗^*_FC_, respectively) in the SP, DP, and random-pattern (RD) cases.

### Testable predictions for simple and complex synapses

Simple and complex synapses exhibit quantitatively different SNR decays and memory performance. We now show that it is possible to design an experiment with a specific learning schedule that would reveal whether the synapses are complex or simple. The main idea is that memories can be repeatedly refreshed in a such way that the asymptotic minimum memory strength remains constant. Ideally, this could be implemented by monitoring the memory signal of a specific stimulus, and refreshing the memory by presenting the same stimulus again as soon as the signal drops below some threshold. Using this procedure, we obtain a refresh schedule, which can be described by specifying the intervals that separate two consecutive presentations of the same stimulus. Depending on the synaptic model, the length of these intervals will be different, and, most importantly, will change over time in a different way.

We illustrate this with three idealized models of memory traces through simulations: the exponential decay model (in which the signal decays as *e^−t/τ^*, where *τ* is the time constant), the inverse-square-root 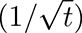 power-law decay, and the hyperbolic (1*/t*) power-law decay (which is achievable by a heterogeneous population of simple synapses^34^). See Fig. 6a-c and f. We set the threshold to *θ* = 0.5, but its precise value does not affect the scaling behavior of the intervals. For the idealized exponential decay model, the length of the interval remains constant after the second presentation (the constant interval is *τ* ln (1 + *C/θ*), where *C* is the initial signal strength and *θ* is the pre-specified threshold). Interestingly, the length of the interval increases asymptotically linearly for the idealized inverse-square-root decay model. The coefficient of this linear asymptotic growth is *π*^2^*C*^2^*/*2*θ*^2^ (see mathematical details in Appendix III). The situation for the hyperbolic decay is intermediate between the exponential and inverse-square-root decay models, showing an approximately logarithmic increase of the length of the interval.

**Figure 6:**
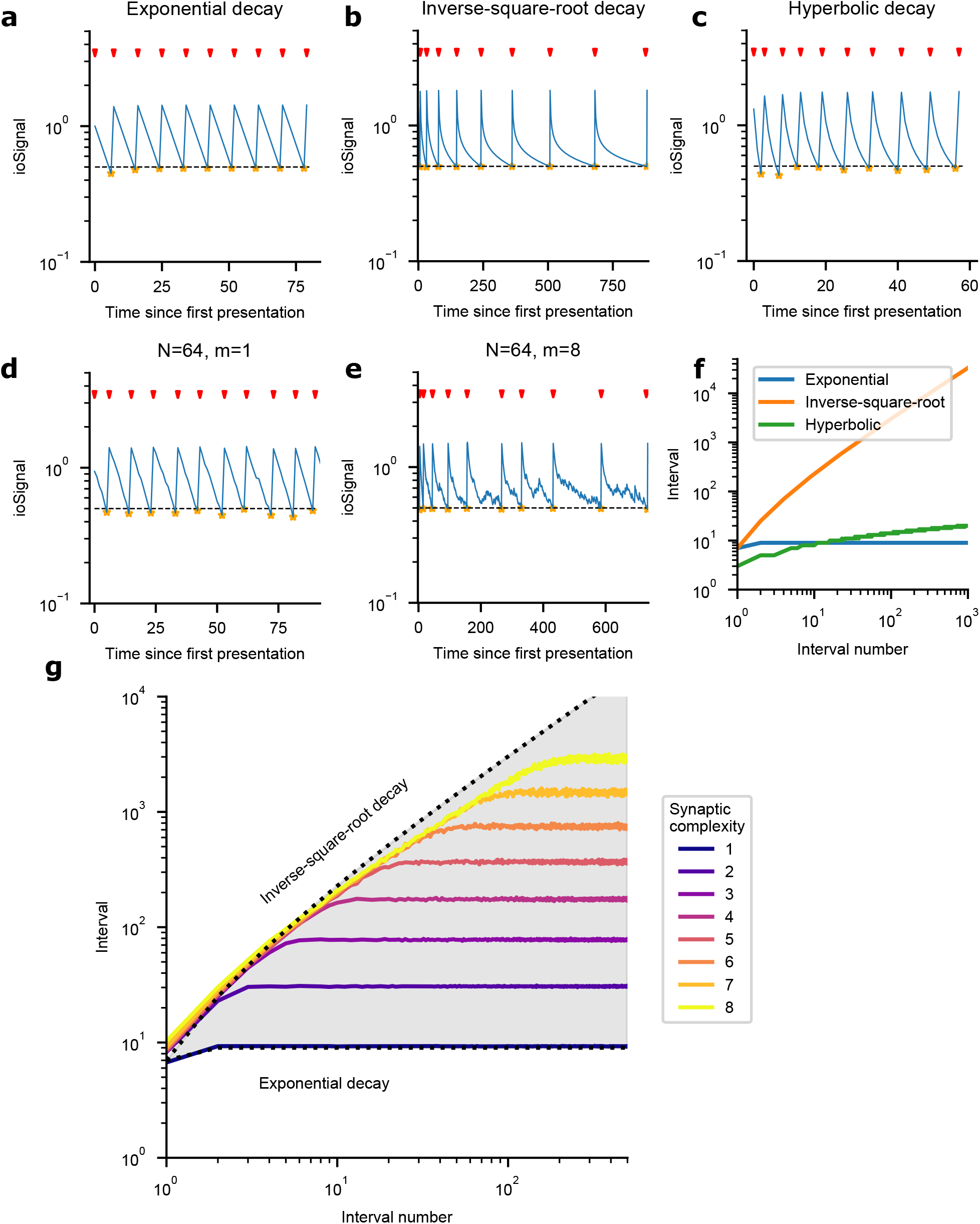
The optimal learning schedule, in which the pattern is represented when the monitored ioSignal drops below a pre-specified threshold (0.5 shown). (a-c) The idealized exponential decay, inverse-square-root decay, hyperbolic decay models of ioSignal under the optimal learning schedule. The ioSignal decreases over time and is enhanced by pattern presentations indicated by red arrows. The signal strength immediately before each presentation is marked by orange stars. (d) A typical noisy ioSignal trajectory from the model with simple synapses (*N* = 64, *m* = 1). The length of the interval between consecutive presentations is approximately constant, similar to the idealized exponential decay. (e) A typical noisy ioSignal trajectory from the model with complex synapses (*N* = 64, *m* = 8). The length of the interval approximately increases linearly with interval numbers, similar to the idealized inverse-square-root decay. (f) The length of the interval as a function of interval numbers for three idealized decay models. (g) The length of the interval increases as a function of interval numbers under the optimal schedule, averaged over noisy ioSignal trajectories. Different colored curves correspond to models with a different synaptic complexity *m*. The shaded region is bounded by interval curves of the idealized exponential decay and inverse-square-root decay models.

The idealized exponential decay and inverse-square-root decay models represent very different degrees of synaptic complexity. The memory signal of complex synapses decays as an inverse-square-root power-law over the longest timescale of the synapse before the decay becomes exponential. Example memory signal trajectories under the optimal learning schedule for simulated models with different synaptic complexity are shown in Fig. 6d-e. The length of the interval is approximately constant for the model with simple synapses (*m* = 1), similar to the idealized exponential decay, but increases linearly for the model with complex synapses (*m* = 8), similar to the idealized inverse-square-root decay. Averaged over multiple such noisy trajectories, the interval curves are plotted on a log-log scale as a function of interval numbers (see Fig. 6g). Increasing synaptic complexity *m* effectively extends the linear growth regime (corresponding to the inverse-square-root power-law decay regime of the ioSNR) and postpones the gradual transition into the constant interval regime (corresponding to the exponential decay regime of the ioSNR). The idealized inverse-square-root decay thus approximates the envelope of the interval curves of models of different synaptic complexity *m*. We further study the scaling properties of the length of the interval in Appendix IV.

However, it would not be feasible in experiments to monitor the memory signal in real time. Indeed, in order to measure the signal we need to expose the subject to the memory we intend to test, and hence we are going to modify the memory signal we want to estimate. We propose that we can simply use either a constant interval or a presentation schedule with a linearly increasing one without monitoring the memory signal (between presentations). Both protocols will be parametrized by a single variable. Under the constant schedule, a specific memory is refreshed after an interval of a fixed length *γ*, while under the linear schedule, a specific memory is refreshed after a linearly increasing interval. In particular, the interval is equal to *γn*, where *n* is the interval number, and *γ* is the length of the first interval (see Fig. 7a,b). These refreshes serve the dual roles of evaluating familiarity detection performance (e.g. by querying the subject whether the presented stimulus is familiar) and boosting memory strength (corresponding to the increase of the signal strength immediately after the refresh). Under the constant learning schedule, the performance of the exponential decay model will remain constant, consistent with the conclusion drawn from the optimal schedule. The inverse-square-root decay and the hyperbolic decay models will exhibit gradually improving performance. Under the linear learning schedule, the performance of exponential decay and hyperbolic decay models will quickly drop to chance level, but the inverse-square-root decay model will maintain its performance, as predicted by the optimal schedule.

**Figure 7:**
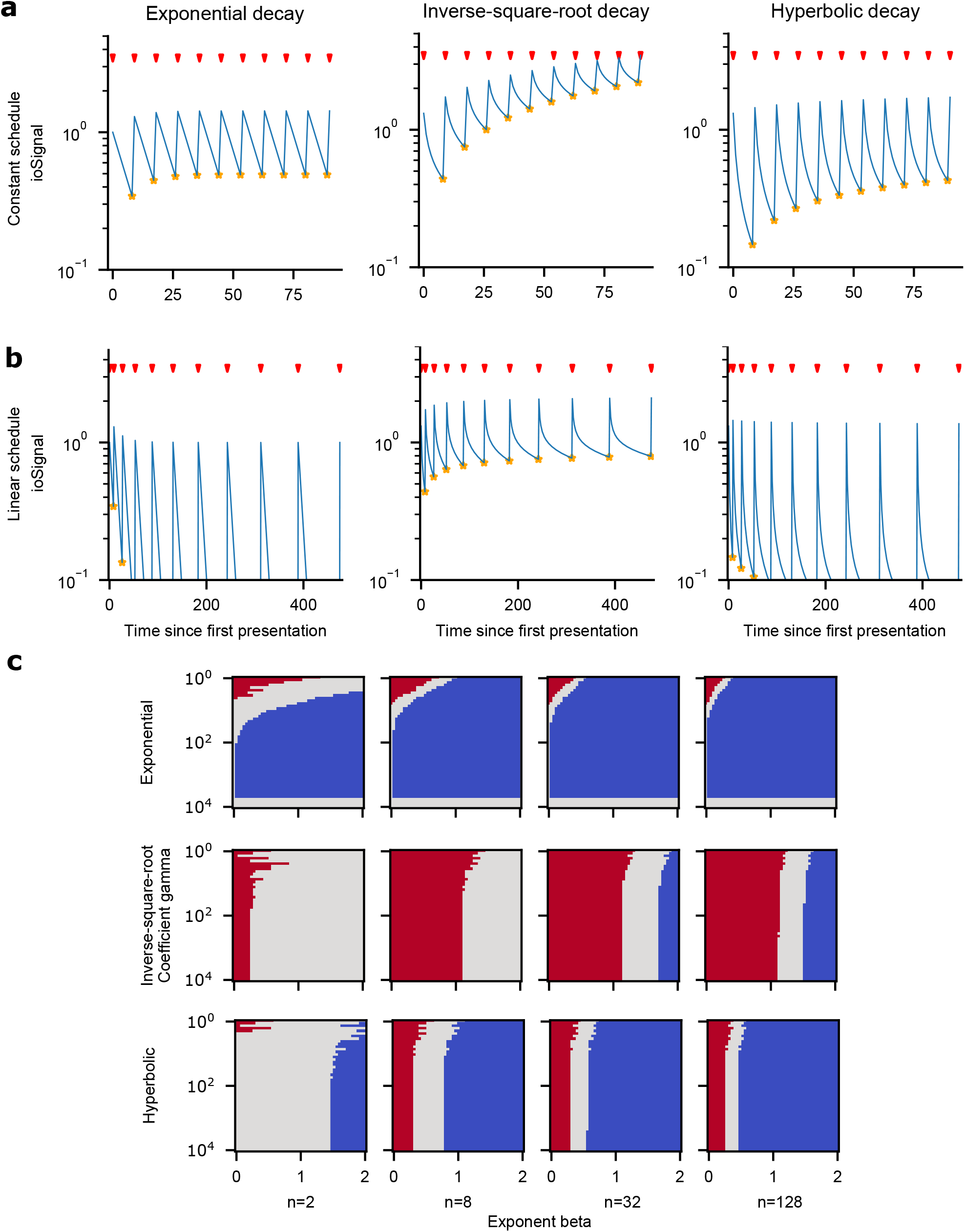
Pre-determined learning schedules. (a) The idealized exponential decay, inverse-square-root decay, hyperbolic decay models of ioSignal under the constant learning schedule, in which the pattern is represented each time after an interval of a pre-determined constant length. The ioSignal decreases over time and is enhanced by pattern presentations indicated by red arrows. The signal strength immediate before each presentation is marked by orange stars, reflecting familiarity task performance. In the following presentations, the task performance of the exponential decay remains constant, while the two power-law decay models’ performance gradually increases. (b) The idealized exponential decay, inverse-square-root decay, hyperbolic decay models of ioSignal under the linear learning schedule, in which the pattern is represented each time after an interval of linearly increasing length. In the following presentations, the inverse-square-root decay model maintains its performance, but the performance of exponential decay and hyperbolic decay models quickly drops to chance level (orange stars not shown due to extremely low signal strength). (c) The signal gain as a function of *γ* (length of the first interval) and *β* (exponent of length increase) for the three idealized decay models under the general pre-determined learning schedule, where the length of the interval equals *γn^β^* (*n* denotes the interval number, 10^0^ *γ* 10^4^ on a log scale, 0 *β* 2 on a linear scale). Red regime is shown for positive gain (*>* 0.2), blue for negative gain (*<* 0.2), and gray for marginal gain (between 0.2 and 0.2). The three idealized decay models exhibit qualitatively different behaviors.

We define the signal gain for interval number *n* as the logarithmic ratio of the ioSignal after the *n*-th interval (immediately before the (*n* + 1)-th presentation) relative to the ioSignal after the first interval (immediately before the second presentation). Positive signal gains correspond to better familiarity detection performance, and negative ones indicate worse performance after the following presentations. The three idealized decay models demonstrate qualitatively different signal gains under the constant and linear schedules with varying *γ* (see Appendix V), offering testable predictions for experiments.

To better discriminate between complex synapses with a square root decay and simple heterogeneous synapses with a hyperbolic decay, we introduced a more general learning schedule (see Fig. 7c). Here the length of the interval takes the form of *γn^β^*, where *n* is the interval number, and *γ* is the length of the first interval. *β* = 0 corresponds to the constant schedule, and *β* = 1 to the linear schedule. In the parameter space spanned by the parameters *γ* and *β*, as the interval number increases, the positive gain regime (red) shrinks quickly for any positive *β* in the idealized exponential decay model. This regime exists in the inverse-square-root decay model for *β >* 1 and in the hyperbolic decay model for smaller but positive *β*. Differentiating between these two power-law decay models requires the examination of the sign of the signal gain around *β* = 1: positive for the inverse-square-root decay and negative for the hyperbolic decay. This general learning schedule thus provides experimental predictions for behavioral signatures that differ between the three idealized decay models, and allow us to discriminate between memory networks of various degrees of complexity.

## Discussion

We have presented a modular memory system that can solve a real-world problem such as face familiarity detection, which involves the ability to store in memory in one shot a large number of visual inputs. Thanks to the interactions between fast and slow variables of the complex synaptic model, the familiarity memory capacity grows almost linearly with the number of plastic synapses or quadratically with the number of neurons. The scaling of the system with simple synapses is only logarithmic with the number of synapses ^3, 33^, though the memory performance can significantly increase when the learning rate *q* becomes small, or when the number of states per variable increases^3, 4^. However, even when the parameter *q* is properly tuned, the linear scaling cannot be achieved with a small number of states, and the system with complex synapses outperforms the one with simple synapses in all cases, even when the total number of dynamical variables is the same for the two systems.

The advantage of complex synapses comes from two important properties: the first one is that they involve multiple timescales, enabling the system to learn quickly using the fast components, and forget slowly due to the slow components. The second one is that the dynamical components operating on different timescales can interact to transfer information from one component to another. In the case of our specific model the information diffuses from the fast components to the slow ones, and back (see^2^ for more details). These two properties are important for any memory system that involves a process of consolidation, whether the process is synaptic or requires communication across multiple brain areas (memory consolidation at the systems level, see e.g. ^34^).

Our previous work^2^ systematically studied the scaling properties, the memory capacity, and the robustness of a broad class of complex synaptic models for random and uncorrelated synaptic modifications. One of the situations in which the synaptic modifications are random and uncorrelated is when the patterns of activity that represent the memories are also random and uncorrelated, which is what was assumed in all the early works on memory capacity (e.g.^22^). One of the reasons behind this assumption is that it allowed theorists to perform analytic calculations. However, it is a reasonable assumption even when more complex memories are considered. Indeed, storage of new memories is likely to exploit similarities with previously stored information. Hence, the information contained in a memory is likely to be pre-processed, so that only those components that are not correlated with previously stored memories are actually stored. In other words, it is more efficient to store only the information that is not already present in our memory. As a consequence, it is not unreasonable to consider memories that are unstructured (random) and do not have any correlations with previously stored information (uncorrelated). Unfortunately, these processes that lead to uncorrelated representations are rarely modeled explicitly (but see^35^) and we currently do not have a general theory for dealing with more realistic, highly structured memories. In our model, the face stimuli, which are highly structured and correlated, are pre-processed by a simulated visual system, whose intermediate representations are then used as inputs to our memory module. This pre-processing seems to be sufficient to achieve approximately the same scaling properties predicted for random patterns.

Another important difference between our previous and present work is related to the nature of the memory problem to be solved. In our previous work, we were dealing either with a classification task with randomly chosen labels (a typical perceptron problem with only one output unit) or with a reconstruction memory problem in which a recurrent network would learn to reproduce a previously seen input at the time of memory retrieval. In this work, we considered familiarity detection, which is a recognition memory problem. To reconstruct each individual binary feature of a memorized pattern, we would employ *N −* 1 synapses. Here we have designed a system in which *N* such output neurons are combined and read out to report a one-bit response, which is familiarity. We are using all *N* (*N −* 1) plastic synapses that are available to output only one bit of information. Hence it is not surprising that in the case of reconstruction memory, the number of memories that can be retrieved (reconstructed) scales linearly with the number of neurons *N*, while in the case of familiarity detection, the memory lifetime scales quadratically with *N* .

We also studied the generalization performance of the system by considering different poses of presented faces as retrieval cues (the DP case), using probe patterns that differ from the originally stored ones. Although the task performance for this DP case is worse than in the SP case, the power-law scaling properties are similar, and the drop in performance could be compensated by introducing more memory neurons and possibly increasing the synaptic complexity. The ability to generalize to different poses is presumably helped by the complexity of the synapses. Indeed, in the case of random patterns, generalization is related to the memory SNR^2^. In future studies, we will determine whether there is a similar relationship between the SNR and the ability to generalize to different poses.

### Biological interpretation

We hypothesize that the embedding module represents the ventral stream of visual cortex, where faces are clearly represented in dedicated patches, which are present in the inferior temporal cortex^36, 37^ and in the perirhinal cortex^38^. The memory module could be mapped onto the hippocampus, containing synapses that can be significantly more plastic than in the cortex. These highly plastic components would support one-shot online learning. This hypothesis would be compatible with the models that see the hippocampus as a memory device that compresses correlated memories before they are stored^39–41^. This compression process is often achieved by modelling the hippocampus as a sparse auto-encoder with one input layer, containing the representation of the memory to be compressed, an intermediate layer and an output reconstruction layer. The weights are tuned to reproduce the input in the output layer. The representations in the intermediate layer are compressed because sparseness is imposed during the learning process. Comparing the input and the output layer would be equivalent to the comparison we perform in our model between the representations in the embedding module and the representations in the memory module. In our model we did not consider an intermediate layer as the face representations are already approximately uncorrelated. However, we could easily introduce an intermediate layer to deal with other classes of visual inputs. The reconstruction layer, and hence the detection module of our model could be in the entorhinal cortex (EC), taking advantage of the architecture of the hippocampal-cortex loop^40^ (the hippocampus projects back to EC, which is also the main input to the hippocampus). Alternatively, it could be that the reconstruction layer is not explicitly implemented (see e.g. ^41^). In this case the compressed representations would emerge in one of the parts of the hippocampus without the need to reconstruct the inputs. It could be in the dentate gyrus, as hypothesized in^41^, or in specific parts of the hippocampus that are involved in social interactions (e.g. CA2 is known to be involved in familiarity detection in mice^42^). The absence of an explicit reconstruction layer would require a more complex readout, that probably needs to be trained because the detection module would have to compare two different representations. This problem could be solved by adopting a different strategy to detect novelty, as suggested in^43^.

Perirhinal cortex is bidirectionally connected with EC and hence with the hippocampus. It could certainly represent familiarity even if we hypothesize that the hippocampus is the main locus of the memory module. This would be compatible with recent electrophysiological observations in monkey perirhinal cortex in which face familiarity is strongly encoded ^38^. This familiarity signal could then be broadcast to the rest of the cortex and explain why familiarity can be decoded also in other areas like infero-temporal cortex^38, 44^.

### Predictions for familiarity detection experiments

Using different learning schedules (including the constant, the linear, and a more general learning schedule), we demonstrate that the exponential decay, the inverse-square-root decay, and the hyperbolic decay models lead to distinct and testable predictions for the familiarity task performance. This can be directly tested in human (and animal) experiments. Within a series of images used in such a familiarity experiment, face images of different identities are ordered so that the same face image is repeatedly presented and at the same time evaluated (by testing familiarity detection) after an interval of a pre-determined length following a specific schedule. How the signal gain (and the corresponding probability of successful familiarity detection) develops as a function of the interval/presentation number will shed light on the temporal decay kernel of the memory signal and therefore on the complexity of the memory consolidation process. We expect that biological synapses will behave similarly to the inverse-square-root model for a wide range of interval numbers until they gradually transition into an exponential decay. If we could ignore the effects of systems-level consolidation and internal replay, the interval number at which this transition occurs would provide a measure of the intrinsic complexity of synapses in the hippocampus.

### Implications for neuromorphic engineering

Because the dynamical variables of the plastic synapses in our memory module are digital and relatively low-precision, they can be effectively implemented as a group of binary switches (bits) in neuromorphic devices, using *b* bits to obtain *M* = 2*^b^* discrete levels. Because the distribution of synaptic weights is approximately Gaussian (though discretized), *M* should be appropriately chosen such that the Gaussian distributions will not be substantially truncated. It is known that for the hidden variables with longer time constants, the width of the distribution is smaller and fewer discrete levels are required^2^. For the hidden variable with the longest time constant, *M* can be as small as 2 without affecting the memory performance much. Another class of complex synaptic models that have been shown to exhibit essentially the same memory performance as the models studied here can be implemented with even fewer hardware bits^45^. Recent developments in the field have shown that memristive devices can be employed in cross-bar architectures to implement synapses with learning capabilities^46, 47^. These devices allow for extremely compact designs as typically they are located at the touch point between axonal and dendritic lines in a cross-bar architecture, effectively allowing for all-to-all connectivity in real-world applications. Adding internal variables with a suitable consolidation dynamics to the plastic synapses, as suggested here, would pose additional challenges for the hardware implementation, but could greatly extend the overall capabilities of the memory system. We believe that the proposed system suggests a useful architecture for a new generation of neuromorphic devices suitable for on-chip online learning. One of the attractive features of the proposed system is that it can solve a real-world problem without requiring a large number of plastic synapses. Indeed, the vast majority of the synapses of the complete system, which would include the pre-processing part, are not plastic. Nevertheless, the system can perform an interesting form of continual learning.

### Limitations of our system

One limitation of our work is the assumption that the memory neurons use exactly the same representations as the input neurons. In reality, the number of memory neurons is unlikely to be precisely the same as the number of input pattern dimensions, and they would in general use a different representation of a given face from the input neurons. The detection module has to essentially compare the reconstructed memory with the representation of the current cue. This is a computation that can be performed even when the representations in the detection module are completely different from those in the input. However, it will require a smarter readout system that is trained to perform this comparison. Generalizing our system to include a more biologically plausible mapping between the embedding module and the memory module, with a corresponding readout mechanism in the detection module, is an important direction for our future work.

In our hippocampus-like memory module, there is only one feedforward layer that uses dense neural representations. However, recurrent neural computations in the hippocampus can be beneficial in some memory tasks^48, 49^. In addition, sparse representations of memory patterns have long been known to harbor computational benefits such as larger memory capacity and the capability to mitigate disruptive effects of correlations ^2, 3, 50, 51^. To what extent recurrent connections and sparse coding are beneficial in our neural system for familiarity detection are questions currently under investigation.

## Author Contributions

All authors conceived the study, participated in the discussions, and wrote the paper. M.K.B. and S.F. supervised the project and acquired funding.

## Acknowledgments

This work was supported by NSF NeuroNex Award DBI-1707398, the Gatsby Charitable Foundation and the Swartz Foundation. M.K.B. was supported by the Kavli Institute for Brain and Mind.

## Data Availability

The VGGFace2 face data set^25^ employed in our study can be found on the following website: http://www.robots.ox.ac.uk/~vgg/data/vgg_face2/.

## Appendix I Choosing the optimal threshold in the FD task

As new patterns are presented to our proposed memory system, the familiarity of any old patterns decreases with time, and therefore the distribution of the readout signal for familiar patterns approaches the one of the unseen patterns (see Fig. 8a). In Fig. 8b, we plot the classification accuracy of the detection module as a function of elapsed time (relative to the presentation of the face pattern) with different signal thresholds. Smaller thresholds result in higher error rates for unseen faces, but have a better performance for familiar faces. To provide an operational definition of familiarity for the detection module, we study the overall performance (the average of classification accuracy over the whole age-range) as a function of the signal threshold and age-range (see Fig. 8c). We include an equal number of familiar and unfamiliar faces in these test sets, and weight false positive and false negative errors equally. For longer age-ranges, the optimal threshold gradually decreases, since the familiar faces become indistinguishable from unseen faces, although choosing the right age-range ultimately depends on the application and on the longest timescale of the synaptic model employed in the memory system.

**Figure 8:**
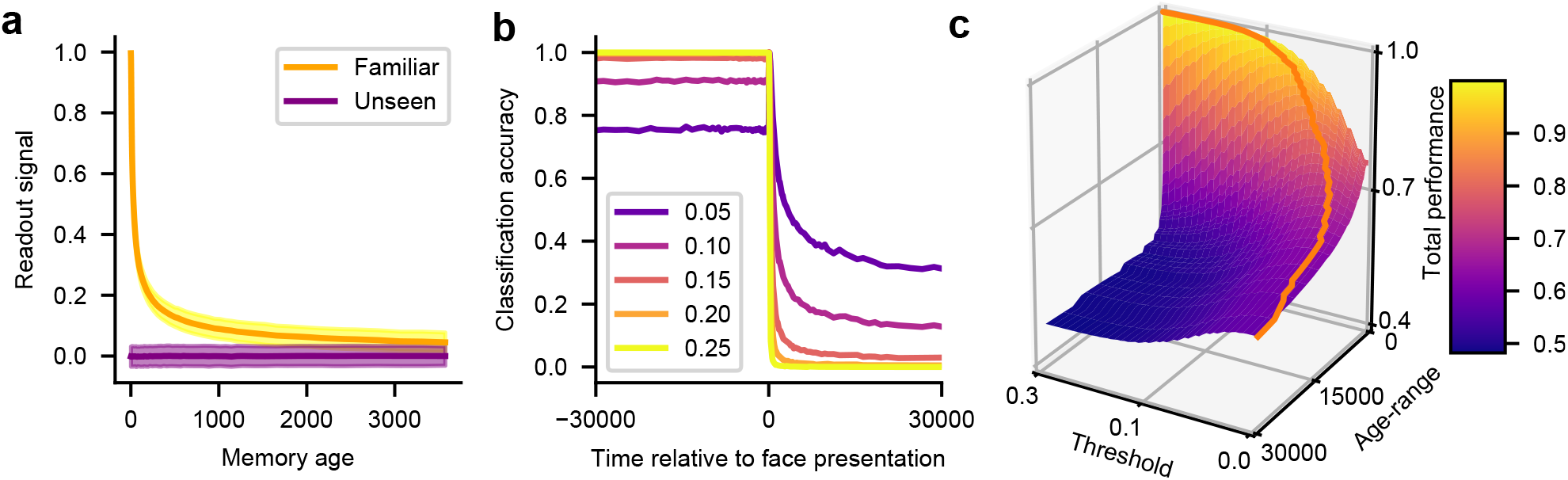
Choosing the optimal threshold. (a) The distribution of the readout signal for familiar patterns approaches the one of the unseen patterns. Shaded regions show 1 SD). (b) The classification accuracy of the detection module as a function of time elapsed since storage with different readout signal thresholds, showing results for the system with *N* = 256 and *m* = 7. (c) The overall performance (the average accuracy over the whole age-range) of the detection module as a function of age-range and signal threshold (again with *N* = 256 and *m* = 7).

For the evaluations in the main text, the threshold of each model is first optimized on a balanced test set with familiar faces within the model-specific age-range (we choose *t^∗^_ioSNR_*), and then evaluated over longer time scales. Since the distribution of the readout signal for unseen faces does not change over time, the detection module with the fixed threshold detects unseen faces with constant error rate (1 - true negative rate) (see Fig. 9b), while it recognizes familiar faces (true positive rate) better for more recent than for older ones (see Fig. 9a). The FD classification performance is the average of the true positive and true negative rates.

**Figure 9:**
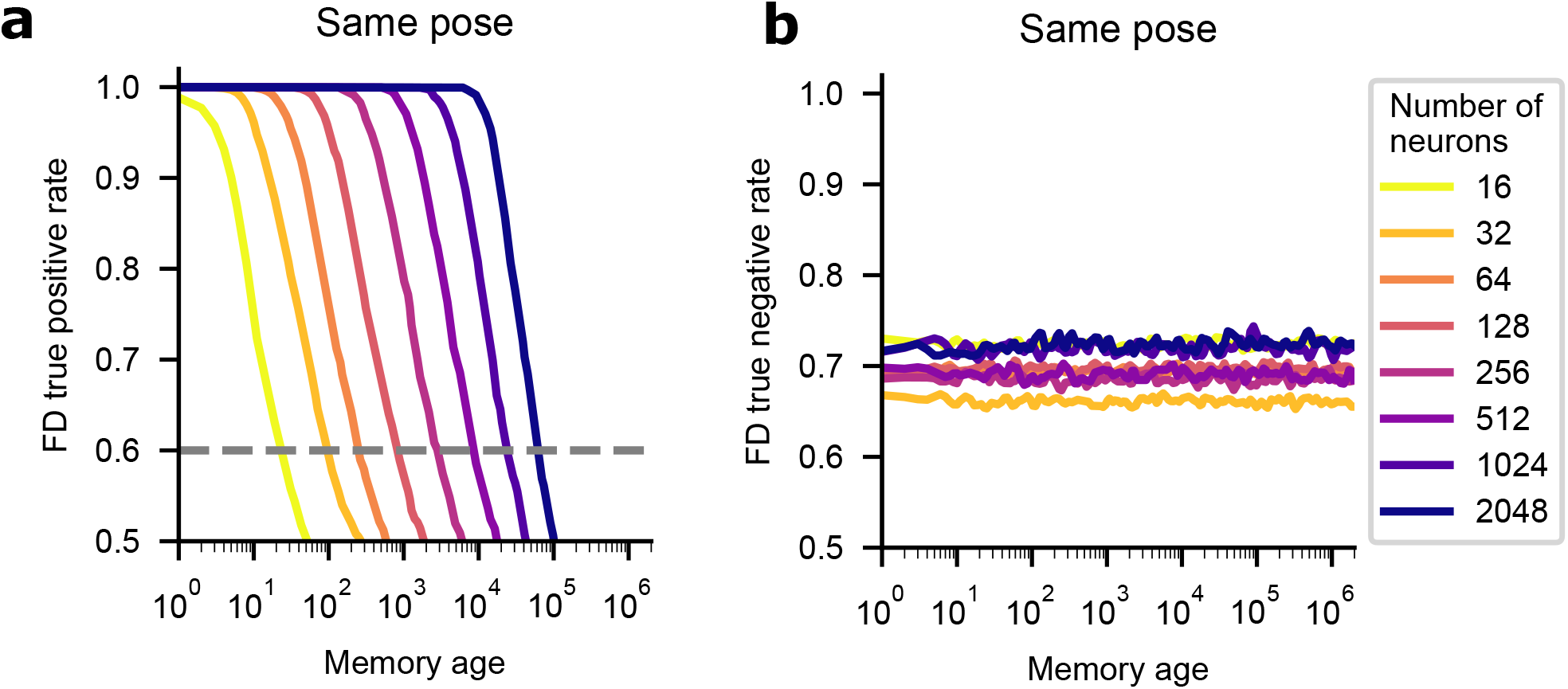
The true positive rate and true negative rate for the FD performance shown in the main text. (a) The true positive rate for familiar faces. (b) The true negative rate for unseen faces, equal to one minus the error rate for unseen faces. Different curves correspond to models with a different number *N* of memory neurons (and number *m* of dynamical variables such that *m* = log_2_ *N −* 1) in the same pose (SP) case.

## II Varying ***N*** and ***m*** separately

In addition to varying the number of memory neurons *N* and the number of dynamical variables *m* together, we also study the properties of our neural system, including the ideal observer signal-to-noise ratio (ioSNR), readout signal-to-noise ratio (rSNR), familiarity detection (FD) test performance, and two-alternative forced-choice (FC) test performance, while varying these two parameters separately.

In the SP case, changing *N* essentially corresponds to shifting the ioSNR curves in the log-log plot vertically (see Fig. 10a for the case with *m* = 8), while increasing *m* corresponds to an elongation of the power-law decay (increasing the longest synaptic timescale *T*) and a slight downward shift of the ioSNR curves within the power-law regime (see Fig. 11b for the case with *N* = 512). With the same threshold as in the main text, the memory lifetime *t^∗^_ioSNR_* scales approximately as *N* ^2^ (see Fig. 10i). Also, the memory lifetime grows exponentially with *m* as long as *N* is sufficiently large (see Fig. 11i). For the DP case (see Fig. 10e, and Fig. 11e), the ioSNR curves are lower than those in the SP case, but still show a reasonable power-law scaling. The rSNR analyses show similar results (Fig. 10b, f, j, and Fig. 11b, f, j).

**Figure 10:**
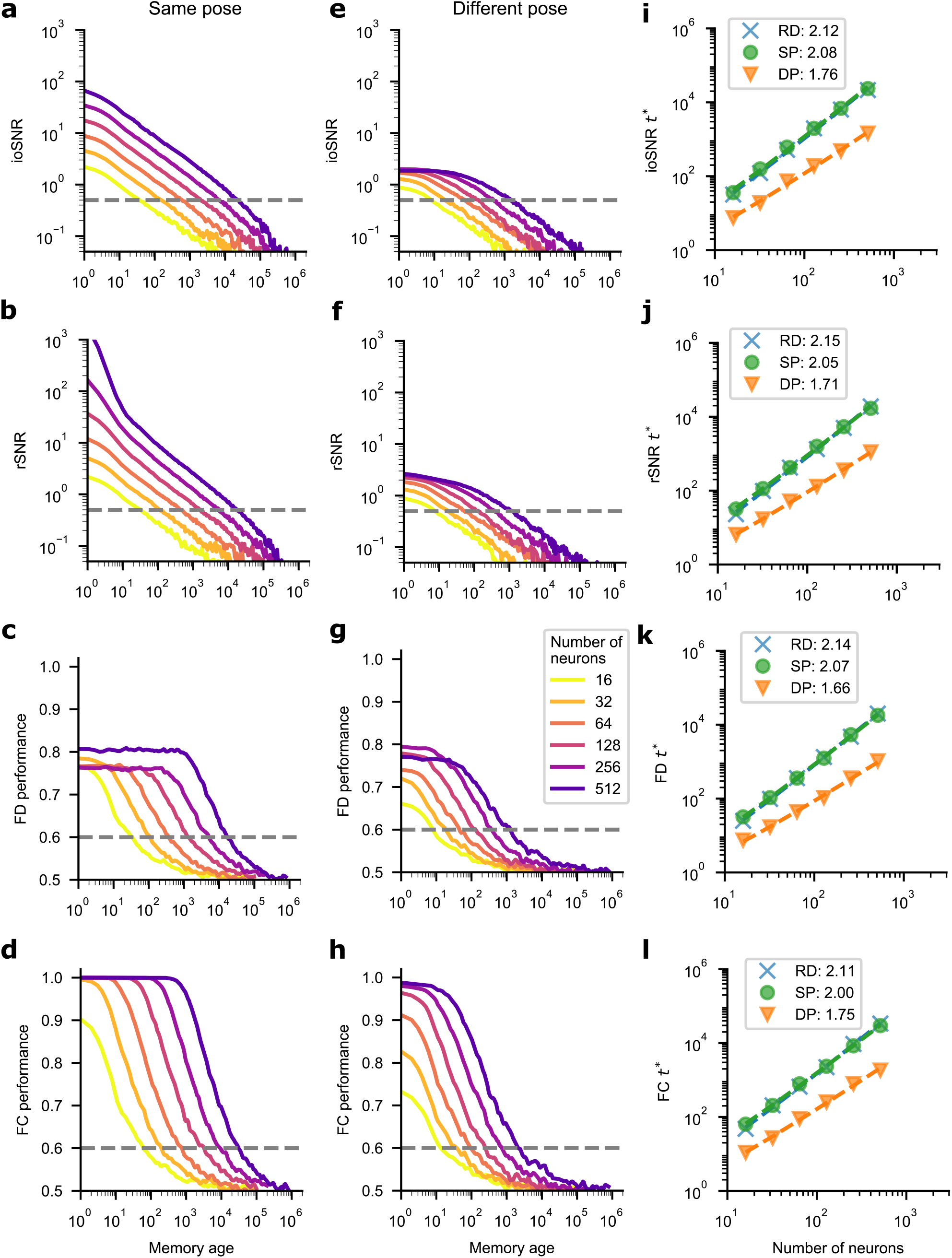
Comparison between models with a different number *N* of memory neurons (*m* = 8 of dynamical variables). (a-d) Comparisons between models for the same pose (SP) case in terms of ioSNR, rSNR, familiarity detection (FD) performance, and two-alternative forced-choice (FC) performance. (e-h) Similar comparisons between models in the different pose (DP) case. (i-l) Comparisons between models in terms of different measures of familiarity memory lifetime (*t^∗^*_ioSNR_, *t^∗^*_rSNR_, *t^∗^*_FD_, and *t^∗^*_FC_, respectively) in the SP, DP, and random-pattern (RD) cases. The legends in (i-l) indicate the best fit linear regression slopes (corresponding to the power of base *N* in the scaling behavior with the logarithmic correction; data points deviating from the regression line due to saturation were excluded).

**Figure 11:**
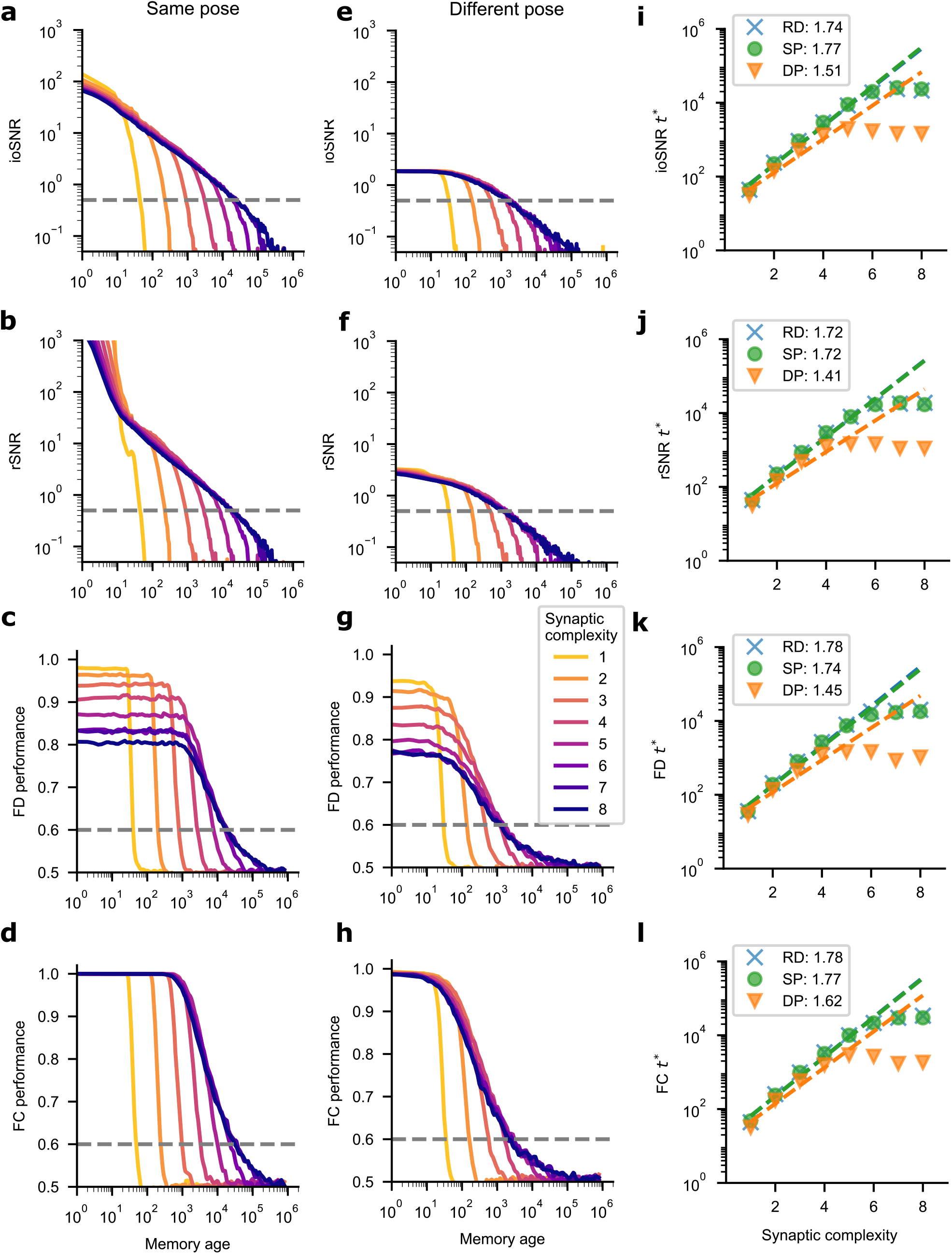
Comparison between models with a different number *m* of dynamical variables (*N* = 512 of memory neurons). (a-d) Comparisons between models for the same pose (SP) case in terms of ioSNR, rSNR, familiarity detection (FD) performance, and two-alternative forced-choice (FC) performance. (e-h) Similar comparisons between models in the different pose (DP) case. (i-l) Comparisons between models in terms of different measures of familiarity memory lifetime (*t^∗^*_ioSNR_, *t^∗^*_rSNR_, *t^∗^*_FD_, and *t^∗^*_FC_, respectively) in the SP, DP, and random-pattern (RD) cases. The legends in (i-l) indicate the best fit linear regression slopes (corresponding to the coefficient of power of base 2 in the scaling behavior with the logarithmic correction; data points deviating from the regression line due to saturation were excluded).

The FD memory lifetime in the SP case is substantially extended by increasing *N* (see Fig. 10c) and *m* (see Fig. 11c). Again, the initial FD test performance cannot reach 100% because each model optimized for a given model-specific age-range constantly make errors for unseen patterns. The FC memory lifetime in the SP case is similarly extended. The initial FC test performance quickly increases as *N* increases in the SP case (see Fig. 10d). Because synapses with fewer dynamical variables have a slightly larger initial SNR (and thus potentially larger performance), the initial performance saturates for models with different *m* between 1 and 8 (see Fig. 11d).

For the DP case, both the FD and FC task performance are worse compared to the SP case (see Fig. 10g, h and 11g, h).

## III Asymptotic behavior of the optimal learning schedules for idealized synaptic models with specific decay kernels

Here we derive the constant length of the interval between successive presentations of the same pattern for a simple synaptic model with an exponential decay and the asymptotically linear increase of the length of the interval for the inverse-square-root decay model.

Let *r*(*t*; *t_n_*) = *r*(*t − t_n_*) denote the decay kernel of the memory signal of a synaptic model at time *t* for a pattern presented at time *t_n_*(the *n*-th presentation, *t ≥ t_n_*). If *θ* is the threshold on this memory signal (such that when the signal has dropped to this level the same pattern will be presented again), we have the following system of equations:

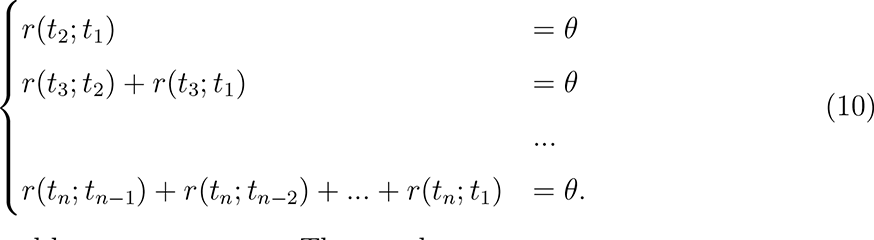

Let the *n*-th interval be *τ_n_* := *t_n_*_+1_ *− t_n_*. Then we have:

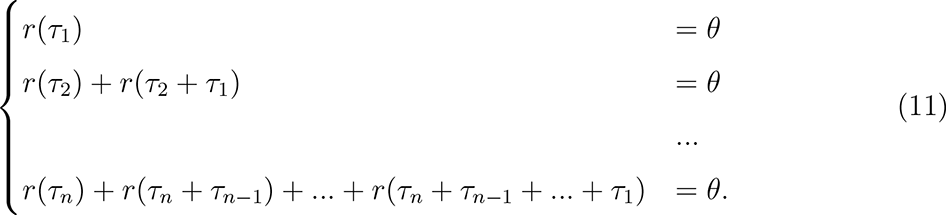

For the exponential decay with *r*(*t*) = *C* exp (*−t/τ*_0_), we can solve these equations from *τ*_1_ to *τ_n_* sequentially and obtain the intervals for the exponential decay:

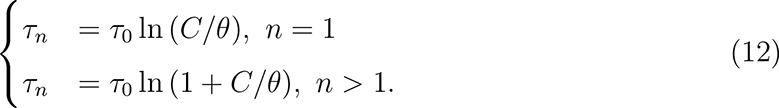

For the inverse-square-root decay with 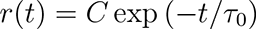, we have:

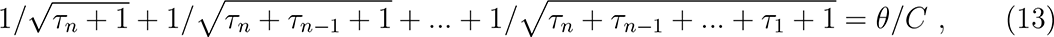

which can be well approximated by the following integral equation when *n* is large enough:

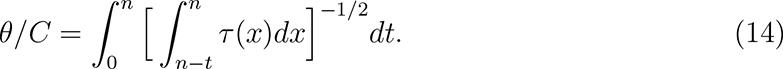

Taking the derivative of both sides with respect to *n*, we have:

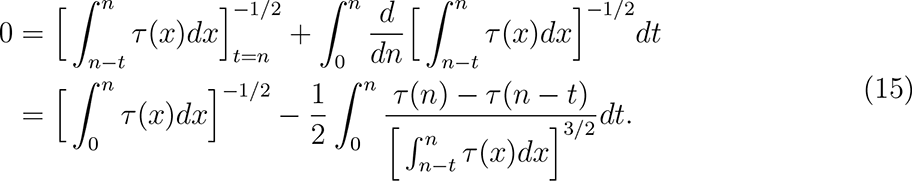

It can be verified that *τ* (*n*) = *Mn* is the solution for the above equation (where *M* is a constant), and thus we proved the asymptotic linearity for the inverse-square-root decay.

To calculate the asymptotic linear coefficient *M*, we insert *τ_n_* = *Mn* into Eq. 13 (and ignore +1 under each square root when *n* is large enough),

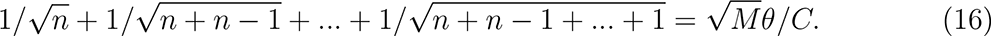

We call the left side *A_n_* and let *n* go to infinity. In this limit we have

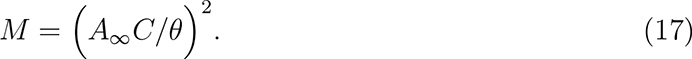

Finally, we calculate *A_∞_* as follows:

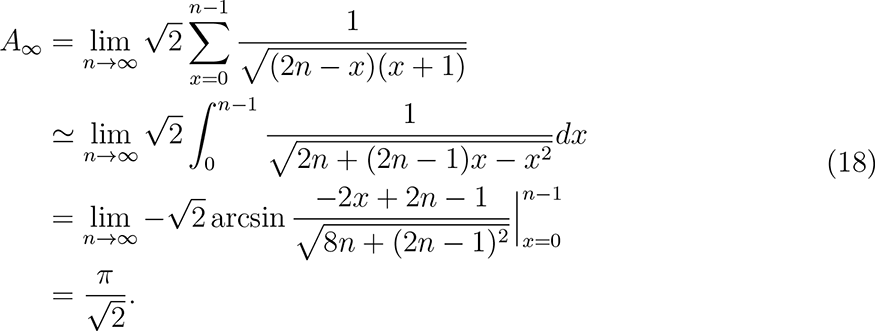

## IV Scaling properties of intervals for complex synapses under the optimal learning schedule

In the main text, we showed that a larger synaptic complexity *m* effectively extends the longest memory lifetime of the synapse and therefore also the linear increase of the length of the interval under the optimal schedule. We computed the limit length of the interval (when the length increase has saturated) and the saturation interval numbers (defined as the time when the interval reaches 99 percent of the limit length) (see Fig. 12a). The limit length of the interval and the saturation interval number scale exponentially with synaptic complexity *m* (Fig. 12b-c).

**Figure 12:**
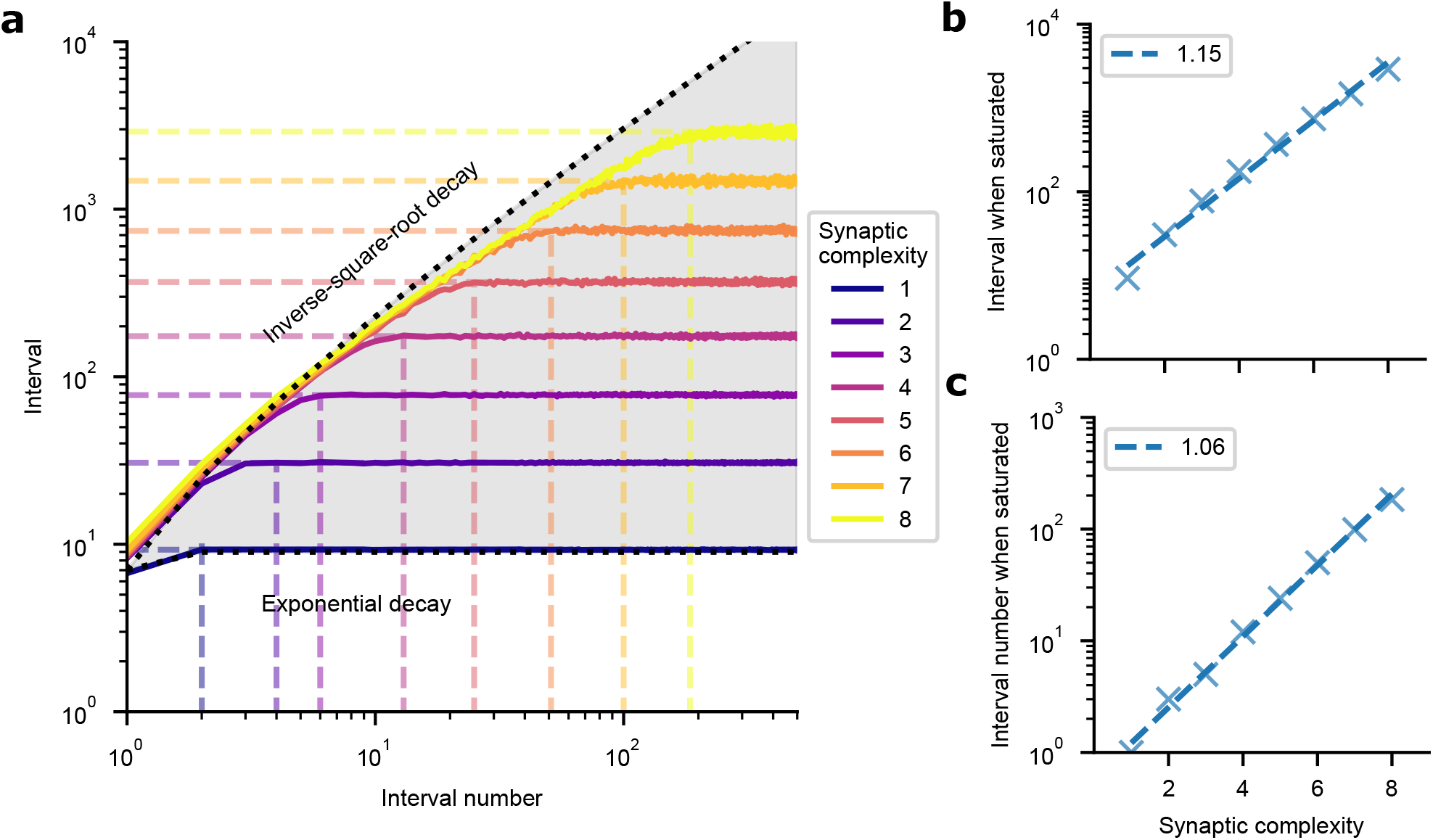
Scaling properties of intervals for complex synapses under the optimal learning schedule. (a) The length of the interval increases as a function of interval numbers under the optimal schedule. Different colored curves correspond to models with a different synaptic complexity *m*. Dotted colored lines correspond to the time when the length of the interval has almost saturated. (b) The limit length of the interval scales exponentially with synaptic complexity *m*. (c) The saturation interval number scales exponentially with synaptic complexity *m*. Labels indicate the regression slope (log_2_ *y ∼ m*).

## V Signal gain under the fixed learning schedules

We studied in detail the signal gain for interval number *n* (the logarithmic ratio of the ioSignal after the *n*-th interval and before the refresh, relative to the ioSignal after the first interval and before the refresh) under the constant and linear learning schedules (see Fig. 13). Positive and negative signal gains correspond to better and worse familiarity performance following subsequent presentations, respectively. The exponential decay model’s signal gain is positive for small *γ* and diminishes for larger *γ* under the constant learning schedule, while it shows a substantial negative signal gain under the linear learning schedule (for *γ* not too small). Under both schedules, the inverse-square-root decay model shows a consistently positive signal gain across all *γ* values. The hyperbolic decay model shows a consistently positive signal gain under the constant schedule, like the inverse-square-root decay model, and a consistently negative signal gain under the linear learning schedule (for *γ* not too small), like the exponential decay model. The differentiation of these three idealized decay models’ performance under both schedules offers testable predictions for experiments.

**Figure 13:**
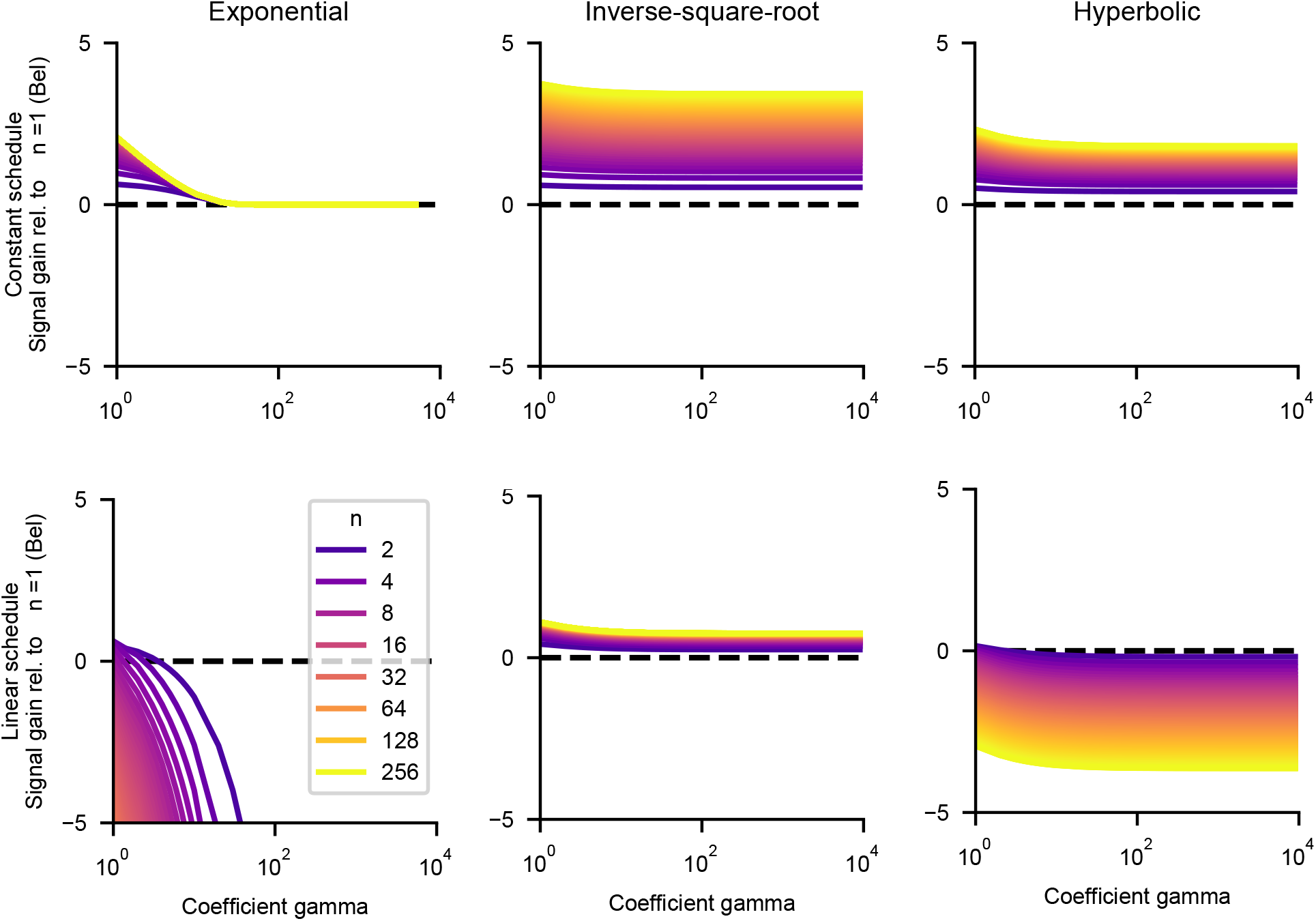
The signal gain as a function of *γ* (length of the first interval in pre-determined learning schedules) for the exponential decay, inverse-square-root decay, and hyperbolic decay models under the constant (first row) and linear (second row) learning schedules. Lighter curves correspond to larger interval numbers.

